# BR-Bodies Facilitate Adaptive Responses and Survival During Copper Stress in *Caulobacter crescentus*

**DOI:** 10.1101/2025.03.11.642215

**Authors:** Christie Passos, Dylan Tomares, Hadi Yassine, Wade Schnorr, Hannah Hunter, Helena K. Wolfe-Feichter, James Velier, Kathryn G. Dzurik, Julia Grillo, Alisa Gega, Sunil Saxena, Jared Schrader, W. Seth Childers

## Abstract

Microbes must rapidly adapt to environmental stresses, including toxic heavy metals like copper, by sensing and mitigating their harmful effects. Here, we demonstrate that the phase separation properties of bacterial ribonucleoprotein bodies (BR-bodies) enhance *Caulobacter crescentus* fitness under copper stress. To uncover the underlying mechanism, we identified two key interactions between copper and the central scaffold of BR-bodies, RNase E. First, biochemical assays and fluorescence microscopy experiments show that reductive chelation of Cu²⁺ leads to cysteine oxidation, driving the transition of BR-bodies into more solid-like condensates. Second, tryptophan fluorescence and EPR assays reveal that RNase E binds Cu²⁺ at histidine sites, creating a protective microenvironment that prevents mismetallation and preserves PNPase activity. More broadly, this example highlights how metal-condensate interactions can regulate condensate material properties and establish specialized chemical environments that safeguard enzyme function.

## Introduction

Biomolecular condensates, which arise through phase separation, are dynamic, membraneless organelles that are critical in coordinating and fine-tuning biochemical pathway specificity. In bacteria, where membrane-bound organelles are largely absent, these biomolecular condensates serve as versatile compartments that organize and regulate key processes such as RNA processing [6, 8], signal transduction [48, 51–53, 59, 60], chromosome segregation [48, 51–53, 59, 60], cell division [54], transcription [55], biofilm regulation [56], and many other processes [1]. One of their defining features is the ability to form reversibly in response to stress, enabling rapid adaptation and recovery [2]. For instance, during heat shock, eukaryotic cells form stress granules that sequester proteins and mRNA, enhancing survival under adverse conditions [3, 4]. For example, yeast species have adapted to diverse thermal environments revealing that while the formation of these condensates is evolutionarily conserved, their properties are fine-tuned to align with each species’ unique ecological niches [5].

Similar stress-responsive condensates are also essential for survival in bacteria. For example, the freshwater bacterium *Caulobacter crescentus* frequently proliferates in environments with limited phosphate. Saurabh *et al.* demonstrated in *Caulobacter crescentus* that phosphate starvation alters ATP levels and, subsequently, the phase behavior of SpmX condensates that regulate cell division [48]. In addition, phosphate depletion also induces BR-body formation, condensates that regulate RNA decay processes in *C. crescentus*, ensuring RNA turnover even when phosphate is limited [6–8]. BR-body formation is further stimulated under various stresses, including ethanol and metal chelator EDTA exposure, suggesting these structures confer adaptive advantages in variable environments [6]. Here, we further explore whether BR-bodies protect bacteria during redox and heavy metal stress, broadening our understanding of biomolecular condensates in microbial stress response.

### BR-bodies as sites of active RNA decay

Our previous studies re-examined the bacterial degradosome in the framework of phase separation and biomolecular condensates [6, 7]. RNase E, an endoribonuclease, performs the critical initial cleavage in RNA degradation and serves as a scaffold via its C-terminal intrinsically disordered region (IDR). This region recruits RNA substrates and downstream degradative enzymes, most notably the 3′-5′ exoribonuclease polynucleotide phosphorylase (PNPase). The degradation process begins with RNase E’s rate-limiting cut, which unfolds RNA, making it accessible and a better substrate for the downstream exoribonuclease PNPase [6, 8–13]. RNase E binds to the helicase RhlB, which enhances degradation by unwinding double-stranded RNA, and also making structured RNA a more suitable ribonuclease substrate [14–16].

RNase E additionally forms biomolecular condensates known as BR-bodies through phase separation. Using RIF-seq [6, 8] and in vitro biochemistry, we showed that recruiting PNPase to BR-bodies enhances mRNA decay [7]. This process is facilitated by RNase E condensates, which accelerate the catalytic activity of PNPase through mass action by bringing the enzymes into proximity to RNA substrates [8]. Notably, *C. crescentus* RNase E selectively binds unstructured RNAs while excluding highly structured rRNAs and tRNAs, tailoring BR-bodies to promote mRNA turnover specifically [6, 7]. After RNase E carries out endonucleolytic cleavage, PNPase and RNase D complete the degradation process through 3′-5′ exonucleolytic activity, converting RNA oligoribonucleotide decay intermediates into nucleotide diphosphates.

The dynamics of BR-bodies are striking, as they form and dissolve rapidly on a sub-minute timescale during mid-log phase growth. The inhibition of RNA polymerase through rifampicin treatment demonstrates that RNA substrates are essential for BR-body assembly [7]. Notably, the C-terminal IDR of RNase E alone is sufficient for phase separation, highlighting the key role of the disordered region in BR-body formation. When untranslated mRNAs form weak multivalent interactions with RNase E, BR-body phase separation is stimulated. This dense liquid-like environment accelerates mRNA degradation and supports a model in which BR-bodies act as dynamic RNA decay centers, assembling and disassembling in concert with RNA degradation (Figure 1A) [6].

**Figure 1:**
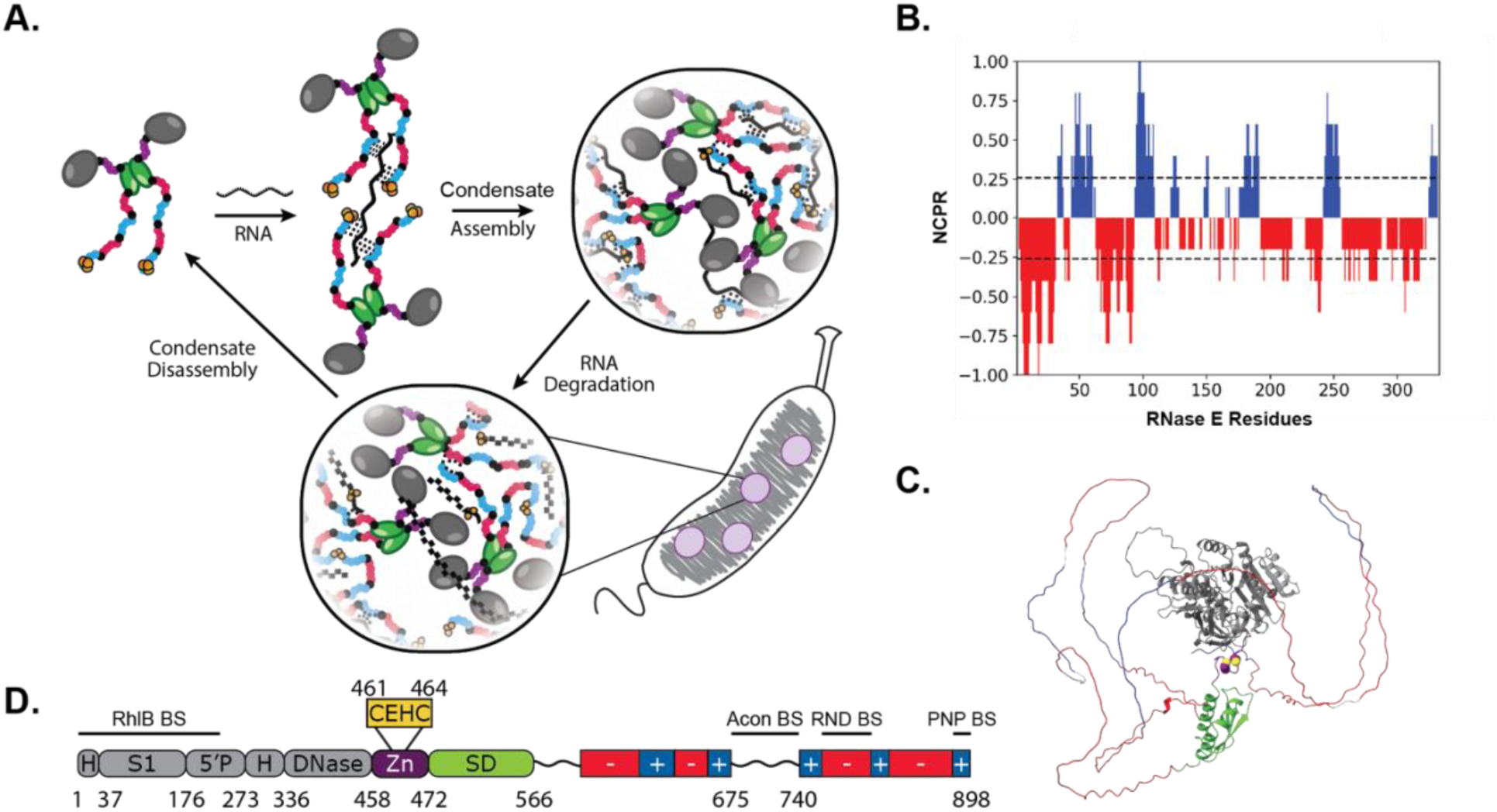
Ribonuclease E (RNase E) is the central scaffold of bacterial ribonucleoprotein bodies (BR-bodies) that mediates global RNA decay in *C. crescentus*. (A) RNase E serves as the central scaffold of the RNA degradosome and is the essential driver of BR-body phase separation. When the degradosome encounters untranslated mRNA, the RNA degradosome makes multivalent interactions with mRNA that triggers phase separation. Mass action effects within the RNase E biomolecular condensate accelerate mRNA degradation and ultimately leads to dissolution of the BR-body upon degradation of the mRNA. (B) A CIDER analysis of RNase E’s IDR reveals a unique linear net charge per residue (NPCR) distribution as discrete alternating charge blocks within the C-terminal domain of RNase E (566-898) [20]. (C) An AlphaFold prediction of *C. crescentus* RNase E (1-898) is shown [21]. (D) Domain architecture of Ribonuclease E. RNase E consists of four domains: the catalytic N-terminal domain (grey, 1-457), the zinc-binding domain (purple, with cysteines highlighted in yellow, 458-471), the dimerizing small domain (green, 472-565), and the disordered C-terminal domain (red and blue, 566-898).

Beyond the core degradosome components, RNase E recruits additional client proteins, such as the Krebs cycle enzyme aconitase, which helps scaffold the RNA decay machinery [13]. Proteomic analyses have identified over 100 RNase E-associated proteins, highlighting BR-bodies’ flexibility in adapting RNA decay to the cell’s changing needs [17]. This interplay between RNase E, RNA substrates, and various client proteins underscores the potential adaptive role of BR-bodies in coordinating RNA decay and maintaining cellular responsiveness to environmental conditions.

### Domains and Functions of Caulobacter crescentus RNase E

Ribonuclease E (RNase E) is a highly conserved bacterial enzyme essential for mRNA turnover. It is present in nearly half of all bacterial species[18]. In *Caulobacter crescentus*, RNase E (CcRNase E) has an architecture comprising a structured N-terminal domain (NTD) responsible for catalysis and an intrinsically disordered C-terminal domain (CTD), which functions as a scaffold and facilitates phase separation. The N-terminal domain (NTD) includes three main regions: the large domain, the Zn-link, and the small domain. The large domain comprises various structural motifs, including an RNase H fold integrated with the S1 RhlB-binding motif, the 5′-phosphate sensors, and a DNase I motif that coordinates catalytic Mg^2+^ ions [19]. Together, these motifs make the NTD chiefly responsible for RNA cleavage and enzymatic activity. The Zn-link and small domain enhance the stability and multimerization of RNase E, while multimerization is further supported by the C-terminal domain (CTD).

The CTD consists of intrinsically disordered regions with alternating positively and negatively charged patches. This arrangement enables the CTD to participate in various weak multivalent homotypic and heterotypic interactions through its interactions with RNA clients. Computational analysis using CIDER the CTD (residues 566–898) highlights its charge distribution, supporting its intrinsic disorder and role in phase separation (Figure 1B).[20]. Moreover, an AlphaFold2 structural model of CcRNase E (residues 1-898) is shown and color-coded to differentiate each region (Figure 1C) [21]. The structured NTD (residues 1-457) appears in grey, the putative Zn-binding domain (residues 458-471) in purple with cysteine residues highlighted in yellow, the dimerizing small domain in green (residues 472-565), and the disordered CTD in alternating red and blue (residues 566-898). In *Escherichia coli*, RNase E’s Zn-link motif stabilizes its oligomeric structure by coordinating a single zinc ion through conserved cysteine residues. It is essential for catalytic activity but is not directly involved in catalysis. Disruption of the Zn-link causes loss of zinc, oligomer destabilization, and inactivation, while artificially maintaining a tetramer preserves activity, emphasizing the motif’s structural role in organizing the active site [22]. In summary, the NTD’s ordered structure supports RNase E’s catalytic role, while the disordered CTD promotes multivalency essential for scaffolding and phase separation (Figure 1D). These domains combine structural order and disorder, enabling RNase E to execute RNA degradation and scaffold assembly for RNA processing machinery efficiently.

#### BR-bodies mediate stress responses

BR-body assembly is not only responsive to RNA substrate availability but also is stimulated by a variety of stresses leading to enhanced viability [6]. When *Caulobacter crescentus* cells were exposed to various acute stresses (e.g., ethanol, EDTA, and heat shock) the stress response included an increased number of RNase E-YFP BR-bodies per cell and an increase in the intensity of the BR-bodies. As previously described by Al-Husini *et al*., under no-stress conditions, an average of 1.1 RNase E-YFP BR-bodies per cell were observed to have formed. In comparison, the presence of 5 mM EDTA for 30 min promoted the formation of 2.0 BR-bodies per cell, while 10% ethanol for 30 min resulted in the formation of 2.1 BR-bodies per cell, and a 5 min 42 °C heat shock produced 2.0 BR-bodies per cell [6]. This past work demonstrates that the C-terminal domain of RNase E is essential for maintaining cellular fitness under a wide variety of stresses.

### Known copper metal response mechanisms in Caulobacter crescentus

*Caulobacter crescentus* thrives in diverse environments due to its ability to withstand various stresses, including those caused by heavy metals like copper [23–26].The bacterium inhabits diverse environments, ranging from freshwater lakes and streams to metal-rich environments like gold mine effluents [27]. Indeed, past studies within *Caulobacter crescentus* NA1000 have identified various heavy metals [23] and oxidative stress response [28] systems.

Copper toxicity in solution arises in part from its high redox potential, which can displace native metallic ions from metalloproteins (leading to mismetallation events) and disrupt Fe–S clusters. Elevated intracellular copper concentrations also trigger a dose-dependent production of ROS, which act as a second messenger in the bacterial stress response, facilitating chemotaxis away from copper [26]. Varying copper stress additionally, modulates the transcriptomic response of both *Caulobacter* cell types to copper stress [23]. The intrinsic dimorphism of *Caulobacter* species with a motile swarmer cell and a non-motile stalked cell allows for a rapid, bimodal response to copper-induced oxidative stress. In stalked cells exposed to elevated copper levels, PcoA oxidizes toxic Cu^+^ to less toxic Cu^2+^, which is then transferred to the PcoB efflux pump in the outer membrane to detoxify the cell. To complement this strategy, *C. crescentus* swarmer cells utilize the cytoplasmic chemoreceptor McpR and the resulting presence of reactive oxygen species (ROS) to mediate chemotaxis away from copper [23–26].

Here, we explore how copper influences BR-body assembly and function, examining the role of phase separation in helping bacteria survive under severe metal stress. We interrogate the functional interactions of copper and RNase E through a combination of in vivo and in vitro assays. These studies collectively point to a model in which BR-bodies facilitate a copper response that enables their survival upon copper stress.

## Results

### CuSO_4_ stress promotes reversible BR-body dissolution in C. crescentus

Previous studies showed that ethanol and EDTA stress enhance BR-body formation [6], prompting us to investigate whether BR-body assembly responds to oxidative and metal stress. Additionally, past studies by Luisi and co-workers have identified that *Escherichia coli* RNase E has a CxxC motif that binds Zn^2+^ ions [22]. Based on these past observations, we investigated the impact of metals and redox stress to determine their effect on BR-body formation (Figure 2A).

**Figure 2:**
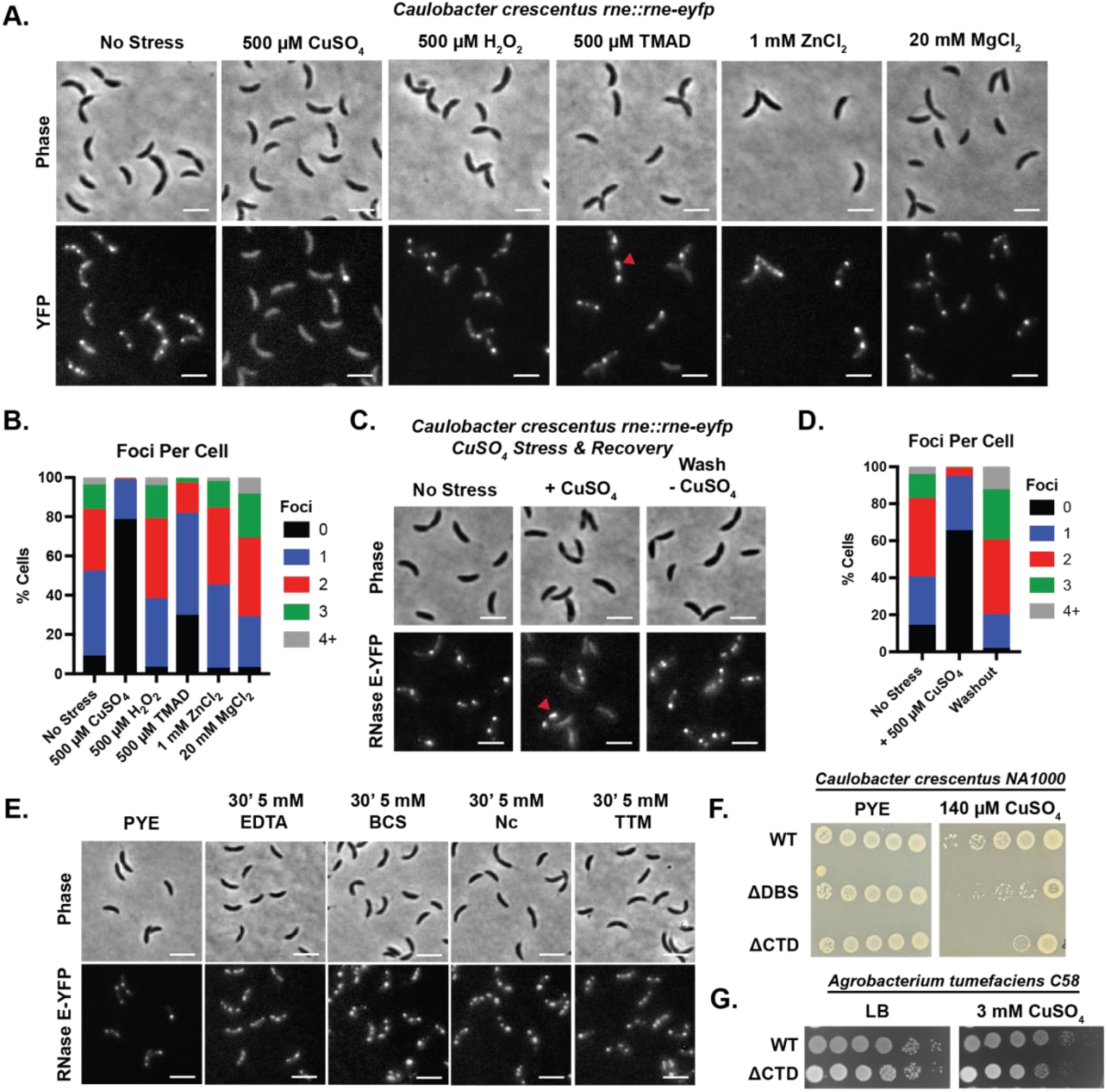
BR-bodies are responsive and provide enhanced fitness when exposed to copper stress. (A) Representative phase contrast and fluorescence microscopy imaging of *C. crescentus* expressing RNase E (1-898)- eYFP from its endogenous promoter (JS51, *rne::rne-eYFP*) in the absence and presence of either 500 µM CuSO_4_, 500 µM H_2_O_2_, 500 µM TMAD, 1 mM ZnCl_2_ or 20 mM MgCl_2_ for 8 min. The scale bar denotes 3 µm. (B) Quantification of the percentage of BR-bodies per cell in *C. crescentus* expressing RNase E-eYFP in the absence (n = 5947) or presence of either 500 µM CuSO_4_ (n = 2289), 500 µM H_2_O_2_ (n = 1613), 500 µM TMAD (n = 1522), 1 mM ZnCl_2_ (n = 957), or 20 mM MgCl_2_ (n = 2285). Amongst stresses, CuSO_4_ selectively leads to dissolution of BR-bodies. (C) Representative phase contrast and fluorescence microscopy imaging of a CuSO_4_ washout experiment with *C. crescentus* expressing RNase E-eYFP from its endogenous promoter (JS51). After CuSO_4_ exposure, cells were washed in fresh medium to remove CuSO_4_ and then re-imaged to observe the recovery of RNase E-eYFP foci after 30 min. We observe recovery of foci after removal of copper from the media. (D) Quantification of the number of BR-bodies per cell pre-CuSO_4_ exposure (n = 870), during CuSO_4_ exposure (n = 1182), and after washout (n = 754). (E) Representative phase contrast and fluorescence microscopy imaging of *C. crescentus* expressing RNase E-eYFP from its endogenous promoter in the absence and presence of 5 mM broad spectrum EDTA metal chelator versus 5 mM specific copper chelators, Bathocuproine sulfonate disodium salt hydrate (BCS), Neocuproine (Nc) and Ammonium Tetrathiomolybdate (TTM) for 30 min. Chelation of metals leads to an increase in BR-body intensity. (F) Recruitment of RNase E clients into BR-bodies and BR-body phase separation provide enhanced fitness under copper stress. Efficiency of Plating assay with wildtype *C. crescentus* RNase E, degradosome binding site mutant (ΔDBS) RNase E and NTD C-terminal deletion mutant (ΔCTD) RNase E in the absence and presence of 130 µM CuSO_4_. Red triangle in panels A and C points to static and elongated BR-bodies in TMAD and CuSO_4_ stress. (G) Efficiency of Plating assay with wildtype A*. tumefaciens C58* RNase E and the RNase e NTD C-terminal deletion mutant (ΔCTD) RNase E in the absence and presence of 3 mM CuSO_4_.

Under no stress conditions, we observed an average of 1.6±0.3 RNase E-eYFP foci/cell in *C. crescentus* cells (Figure 2B, S1B, S1D, S1F). In comparison, when cells were exposed to CuSO_4_ stress for 8 min (Figure 2A), cells displayed a more diffuse subcellular localization pattern with considerably fewer BR-bodies by 500 µM CuSO_4_ with an average of 0.24±0.5 foci/cell (Figure 2A, 2B, S1A, S1B). Notably, about 20% of cells contain BR-bodies that were larger and more intense than the no stress condition. We next considered whether this CuSO_4_-mediated reduction in foci resulted from oxidative stress. We found that H_2_O_2_ exposure for 8 min had little impact on the subcellular localization, with an average of 1.85±0.9 foci/cell (Figure 2A, 2B, S1F, S1G). In comparison, adding 500 µM TMAD led to a decrease to 0.92±0.8 foci/cell (Figure 2A, 2B, S1D). Additionally, a small population of cells exposed to TMAD formed larger and brighter BR-bodies than those observed in the no stress condition. However, increasing amounts of TMAD from 600 µM to 1 mM could not fully dissolve the condensates, as we observed with CuSO_4_ addition (Figure S1D, S1E). This suggests that the dissolution of BR-bodies by CuSO_4_ may be a specific copper effect, as opposed to a generalized oxidative stress effect.

Given that BR-body assembly is enhanced upon EDTA exposure (Figure 2E, S2A), we considered the impact of other metals commonly chelated by EDTA, including Mg^2+^ and Zn^2+^ [6]. Unlike CuSO_4_, adding 1 mM ZnCl_2_ for 8 min had little to no impact on RNase E’s subcellular localization, with 1.7±0.9 RNase E-eYFP foci/cell (Figure 2A, 2B). In comparison adding 20 mM MgCl_2_ for 8 min led to a moderate increase in BR-bodies to 2.1±1.1 RNase E-eYFP foci/cell (Figure 2A, 2B). This impact of magnesium is consistent with our past in vitro observations that magnesium ions could stimulate RNase E phase separation in vitro[29]. So, while the addition of MgCl_2_ results in an increase on BR-bodies, CuSO_4_ has a distinct effect of mediating substantial dissolution of BR-bodies.

Next, we considered that CuSO_4_ might have a nonspecific effect on the subcellular localization of other biomolecular condensates. Thus, we examined an additional protein that phase separates in *C. crescentus*, PopZ, a cell-cycle signaling scaffolding protein that localizes to the cell poles. Unlike the impact on RNase E’s subcellular localization, we found that 500 µM CuSO_4_ did not affect the subcellular localization of PopZ-mCherry condensates (Figure S1C). This suggests that CuSO_4_ specifically impacts BR-bodies directly or indirectly.

Biomolecular condensates are characterized by their capacity to undergo rapid and reversible changes in formation and dissolution upon changes in stress [2]. We thus considered whether BR body formation would recover after removing CuSO_4_ stress (Figure 2C, 2D). Before adding copper, we observed 1.69±1.1 foci/cell, but after exposure to 500 µM CuSO_4_ for 8 min, the number of foci per cell decreased to 0.41±0.6.

Interestingly, amongst the 33.9% that display one or two foci, there exists a population of cells with larger and more elongated BR-bodies than those observed in the no stress condition (Figure 2C, red triangle). After washing cells in fresh medium for over 30 minutes to remove CuSO_4_, we observed complete recovery of BR-bodies, with cells displaying 2.39±1.2 foci/cell (Figure 2C, 2D). Interestingly, analysis of average total cell intensity revealed a 54.4% decrease upon addition of CuSO_4_ (Figure S1H). This suggests that copper binding to RNase E may cause localized quenching of YFP, potentially reducing our ability to detect BR-bodies. Overall, our findings demonstrate that CuSO₄ stress induces a reversible and RNase E-specific effect on RNase E subcellular localization in C. crescentus cells.

Having previously observed the ability of CuSO_4_ to dissolve BR-bodies at short times (8 min) (Figure 2A, 2B), we examined the impact of CuSO_4_ at an extended 1-hour exposure (Figure S2C). After a 1-hour exposure to CuSO₄, *C. crescentus* cells exhibited an increase in RNase E-eYFP total cell intensity (Figure S2D). This suggests that prolonged CuSO₄ exposure may influence RNase E protein expression levels. Notably, the 5ʹ-UTR of the *rne* transcript is a substrate of BR-bodies [7], and copper-induced changes in BR-body phase separation could alter RNA half-life, potentially affecting protein expression. Additionally, extended exposure may allow cellular responses to mitigate cytoplasmic copper levels through reduction mechanisms [23–26].

### Copper-specific chelators lead to increased BR-body intensity

Given the acute impact of copper exposure on the dissolution of BR-bodies (Figure 2A, 2B) and the increase in BR-body intensity when exposed to EDTA (Figure 2E, S2A), we wondered if these phenomena are connected. EDTA exhibits a strong preference for chelating metal ions, including calcium (Ca^2+^), magnesium (Mg^2+^), and copper (Cu^2+^), along with other transition metals such as iron (Fe^2+^/Fe^3+^), lead (Pb^2+^), and zinc (Zn^2+^). (Figure S2A) [45]. We observed that EDTA treatment led to an increase in BR-body intensity, rising from 972 ± 257.81 arbitrary units (AU) (no stress) (n = 2803) to 1297 ± 291.45 AU (n = 1272), (*p* < 0.001), (Figure S2A), consistent with our previous findings [6].

To test if more specific metal chelation could be sufficient, we compared the effects of EDTA with three copper-specific chelators: bathocuproine sulfonate (BCS), neocuproine (Nc), and ammonium tetrathiomolybdate (TTM) (Figure 2E). BCS preferentially binds Cu^1+,^ while TTM selectively binds Cu^2+^ (Figure S2A). In comparison, Nc more promiscuously interacts with various metals, including Cu^1+^, Fe^2+^, and Zn^2+^ (Figure S2A). We found that these copper-specific chelators increased BR-body intensity compared to the no-stress condition. In the absence of stress, we measured an intensity of 972 ± 257.81 AU (n = 2803), while the intensities for the copper chelators were 1383 ± 370.75 AU (BCS, *p* < 0.001) (n = 1876), 1544 ± 310.63 AU (TTM, *p* < 0.001) (n = 1775), and 1337 ± 297.92 AU (Nc, *p* < 0.001) (n = 2265) (Figure S2A).

Since copper-specific chelators increase BR-body intensity, we wondered whether the BR-bodies could sequester copper under conditions with low metal content. Such a material response would share similarities to the observation of enhanced BR-bodies during phosphate nutrient limitations [8]. To test this hypothesis, we examined whether RNase E’s ability to recruit clients and undergo phase separation enhances fitness during metal chelation (Figure S2B). We performed an efficiency of plating (EOP) assay to compare the growth of wild-type *C. crescentus* with strains expressing either the RNase E N-terminal domain (NTD) alone or a mutant lacking the C-terminal domain (ΔCTD) under copper-specific chelation. The EOP results (Figure S2B) indicate that when cells are grown under EDTA chelation, Nc chelation, or TTM chelation, strains lacking the capacity to form BR-bodies demonstrated reduced fitness. In comparison, cells expressing full-length RNase E or the Rnase E degradosome binding site mutant displayed robust growth in the presence of the tested chelators. These findings suggest that under metal-depleted conditions, the RNase E condensate environment may help facilitate client binding to essential metal cofactors. Additionally, prolonged metal depletion stress on solid media may induce changes in *RNase E* transcription that promote survival.

### BR-body formation provides high fitness during copper-induced stress

Given the impact of copper stress on the subcellular localization of RNase E-eYFP, we investigated whether these biochemical processes are critical for cell fitness. To address this, we examined the capacity of *C. crescentus* to survive copper stress using an EOP assay (Figure 2F, S3B). We observed colony growth in wild-type strains at CuSO_4_ concentrations as high as 200 µM (Figure S3B). This level of copper toxicity aligns with findings from Matroule and colleagues, who reported a substantial decrease in the cumulative mass of *C. crescentus* [24] under similar CuSO_4_ stress conditions in PYE [24]. Next, we explored if RNase E’s phase separation capacity and client recruitment functions contributes to copper stress response in *C. crescentus*. We constructed a *C. crescentus* strain that expressed only the N-terminal domain (NTD, residues 1-577) of RNase E, which neither phase separates nor recruits client proteins (JS769). To differentiate the effects of client recruitment from phase separation, we utilized a strain with deletions of the binding sites for PNPase, aconitase, and RNase D (RNase E ΔDBS, JS801). Previous studies have shown that the RNase E ΔDBS mutant can phase separate into BR-bodies but not recruit PNPase, RNase D, or aconitase [6].

EOP assays conducted in the presence of 130 µM CuSO_4_ revealed that wild-type strains exhibit enhanced fitness compared to RNase E ΔDBS strains (Figure 2F). Similarly, we examined the fitness of *Agrobacterium tumefaciens* in LB supplemented with 3 mM CuSO_4_, comparing the wild-type strain to a strain expressing RNase E ΔCTD as the sole variant (Figure 2G, S3C). We found that *Agrobacterium* with RNase E is incapable of stimulating phase separation and exhibited reduced cell fitness. These results collectively suggest that the protective role of the BR-body phase-separated environment is also conserved in *Agrobacterium* and recruiting PNPase, aconitase, RNase D, and potentially other clients into BR-bodies could be critical for maintaining high fitness under copper stress. This may be because BR-bodies provide a protective zone that retains the functional capabilities of client proteins. Alternatively, the client proteins themselves may play some role in attenuating the toxicity of the copper metals. Additionally, removal of the client binding sites may impact the recruitment of the other 111 client proteins of BR-bodies [50].

In contrast, the RNase E NTD variant, which also lacks phase separation capability, demonstrated even lower fitness in the presence of CuSO_4_. This indicates that BR-body phase separation capacity plays a significant role in *C. crescentus*’ response and improved fitness during copper stress. Interestingly, despite copper’s general toxicity to *C. crescentus*, in the presence of copper, the mRNA half-life changes little upon copper exposure (i.e., 0.52 min, while at 125 µM CuSO_4,_ the half-life was 0.79 min, and at 500 µM, the half-life was 0.46 min, as shown in Fig S3D and S3E). This supports the idea that the degradosome function is insensitive to CuSO_4_ up to 125 µM.

### The CxxC motif contains sequence variation in the flanking residues

Given the observed effects of CuSO_4_ on BR-body formation and its role in supporting high fitness during copper stress, we aimed to investigate if the conserved CxxC motif in the annotated Zn-link was involved in zinc or copper binding. Past biochemical studies of *E. coli* RNase E provide evidence for a conserved CxxC motif that mediates tetramerization in a Zn^2+^-dependent manner of *E. coli* RNase E [22]. The CxxC motif is well-conserved across the α-proteobacteria (CPHC, Figure 3A) and in the γ-proteobacteria (CPRC). The consensus sequences of the α-proteobacteria (CPHC) and γ-proteobacteria (CPRC) are highly similar, both containing a proline in the second position. Proline induces β-turns and can specifically impart rigidity to CxxC motifs if placed in the second or third position [49]. Uniquely, *C. crescentus* RNase E has lost that proline in its second position (CEHC, Figure 3A), which could suggest it is more flexible. The next neighboring amino acid of *E. coli* RNase E (CPRCS) also differs from that of *C. crescentus* (CEHCE). Overall, the introduction of glutamates and the replacement of arginine for histidine may alter metal-binding preference. Additionally, the CxxC motif is also known to oxidize to generate disulfide bonds. The bacterial thioredoxin *Bascillus subtilis* ResA utilizes a CxxC motif in a loop region that can form disulfides. Therefore, the *C. crescentus* RNase E CxxC motif may have modified metal-binding preferences or be more readily oxidized to form disulfide bonds.

**Figure 3:**
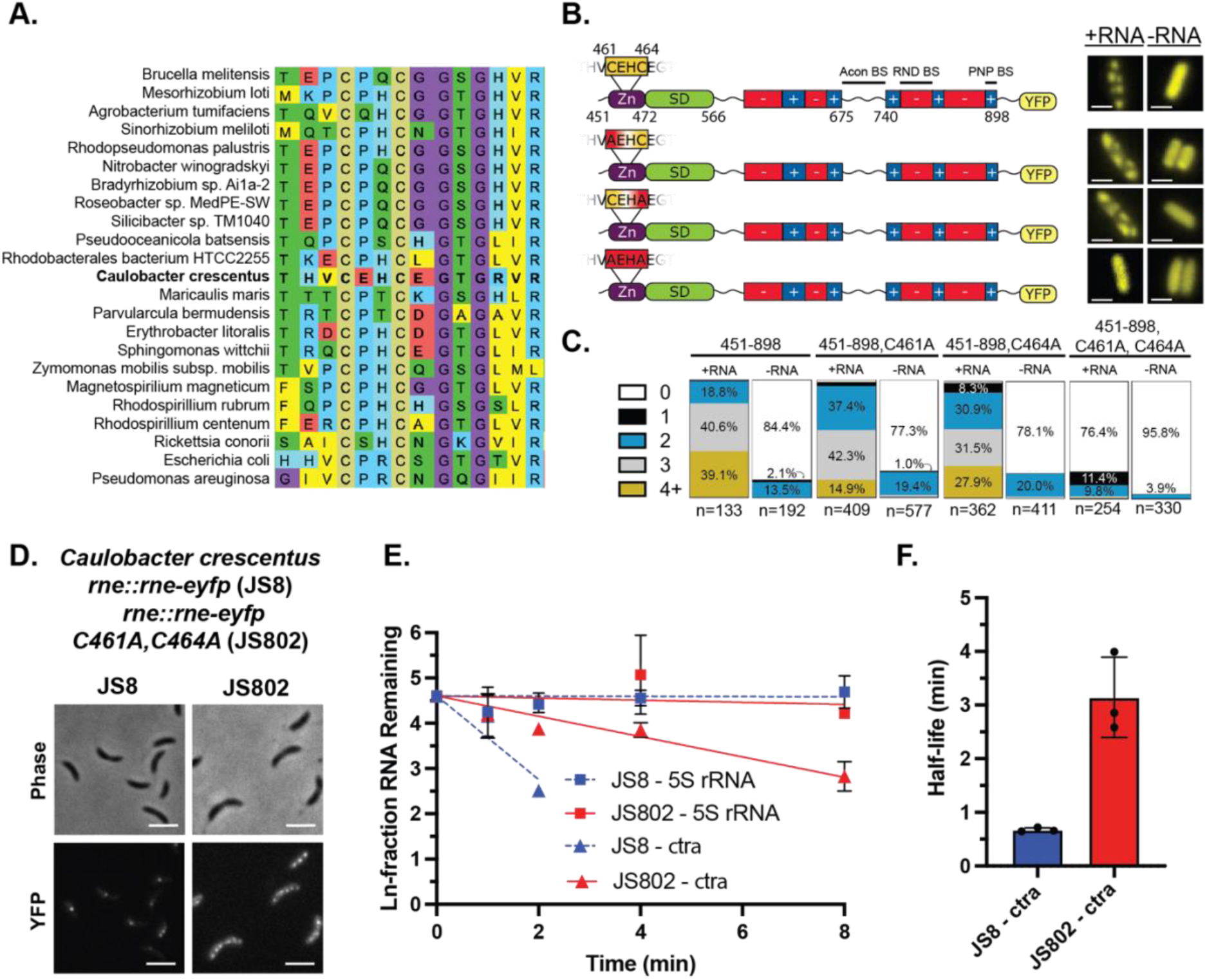
RNase E CxxC motif plays a critical role in BR-body assembly and ribonuclease function in vivo. (A) The CxxC motif is conserved throughout α-proteobacterial RNase E homologs. (B) Comparison of RNase E(451-898)-eYFP, RNase E(451-8987, C461A)-eYFP, RNase E(451-898, C464A)-eYFP and RNase E(451-898, C464A,C461A)-eYFP when heterologously expressed in *E. coli* demonstrates that at least one cysteine is necessary for foci formation. (C) The quantification of foci in the panel above supports an increase in the number of foci/cell when at least one cysteine is present. (D) Representative phase contrast and fluorescence microscopy imaging of *C. crescentus* expressing RNase E (451-898)-eYFP from its endogenous promoter (JS8) and *C. crescentus* RNase E(451-898), C461A, C464A)-eYFP variant. (E) Comparison of RNA half-life for 5S rRNA and ctrA for wildtype RNase E vs. mutant RNase E CTD C464A C461A. (F) Quantification of mRNA half-life for CtrA for wildtype RNase E vs. mutant RNase E CTD C464A C461A.

### C. crescentus RNase E C461 and C464 contribute to phase separation in vivo

First, we considered the importance of the cysteines to phase separation of RNase E in vivo through heterologous expression of RNase E variants designed as single and double mutations of the conserved cysteines (C461 and C464) to alanine (Figure 3B) within C-terminal domain (residues 451-898) which is sufficient for phase separation of RNase E [6]. A single mutation of either cysteine (C461 or C464) to alanine did not abolish phase separation (C461A: 98.0% localized, n=409; C464A: 98.6% localized, n=362), and cells displayed RNA-responsive foci. However, the double cysteine to alanine mutant displayed a 76.4% diffuse protein population (n=254), suggesting that functional cysteines are necessary for RNase E(451-898)-eYFP foci formation in *E. coli* (Figure 3B, 3C).

Our findings that at least one cysteine is required for the phase separation of heterologously expressed CcRNase E in *E. coli* (Figure 3B) prompted us to explore whether this also applies to CcRNase E when expressed in *C. crescentus*. To test this, we generated a CcRNase E C461A/C464A variant lacking both cysteines and assessed its ability to form foci (JS802) (Figure 3D). Interestingly, the results show that this variant can still form foci and even displays an increased average number of foci per cell compared to the *rne::rne-eyfp* control (Figure S3A). Specifically, the wild-type RNase E-eyfp displays an average of 1.2±0.6 foci/cell (n = 473), while the C461A/C464A variant shows 1.8±1.3 foci/cell (n=354), a 28% increase (Figure S3A). Notably, as the C461A/C464A variant is expressed, the unlabeled wild-type is depleted in this strain. So, a small amount of wild-type RNase E may influence the subcellular localization of the C461A/C464A variant.

When examining partition ratios (the average focus intensity over background), the wild-type control has a significantly higher average partition ratio (4.24 ± 0.28, n = 473) than the C461A/C464A variant (2.81 ± 0.30, n = 354) (Figure S3A). This suggests that the absence of cysteines may lead to more diffuse or less tightly packed protein localization. The increased localization of RNase E in *C. crescentus* could be influenced by the inclusion of the N-terminal domain, which may enhance valency, cognate RNAs present in the cellular milleu, or by specific clients in *C. crescentus* that promote phase separation of the C461A/C464A variant. Nonetheless, the data indicates that the RNase E C461A/C464A variant exhibits altered subcellular localization. In addition, we observed that in the presence of the RNase E C461A/C464A variant, the ctrA mRNA half-life was substantially increased to 3.14 min versus 0.67 min for wild-type (Figure 3E and 3F). This is consistent with variants’ impact on the phase separation functions of the CTD, as well as the roles of the cysteines in the RNase E’s ribonuclease domain reported by Luisi and co-workers [21].

### RNase E’s cysteines mediate reductive chelation of Cu^2+^ to Cu^1+^

Previous studies by Luisi and colleagues identified that *E. coli* RNase E contains a CxxC motif that binds Zn^2+^ ions and mediates tetramerization [22]. We, therefore, used the Ellman assay to measure the number of free thiol groups in the presence and absence of various metals to assess metal-binding preferences (Figure 4A). The assay revealed that zinc did not significantly reduce the free thiol content compared to reduced RNase E (451-898) (2.0±0.1, *p* = 0.3390), suggesting that Zn^2+^ does not bind to RNase E (451-898). Significant thiol reduction was observed only with Cu^2+^ (0.1±0.1, *p* < 0.0001) and Cu^1+^ (0.1±0.1, *p* < 0.0001; all others *p* ≥ 0.3390) (Figure 4A), indicating that the zinc-binding domain functions as a Cu-mediated sensor in *C. crescentus*.

**Figure 4:**
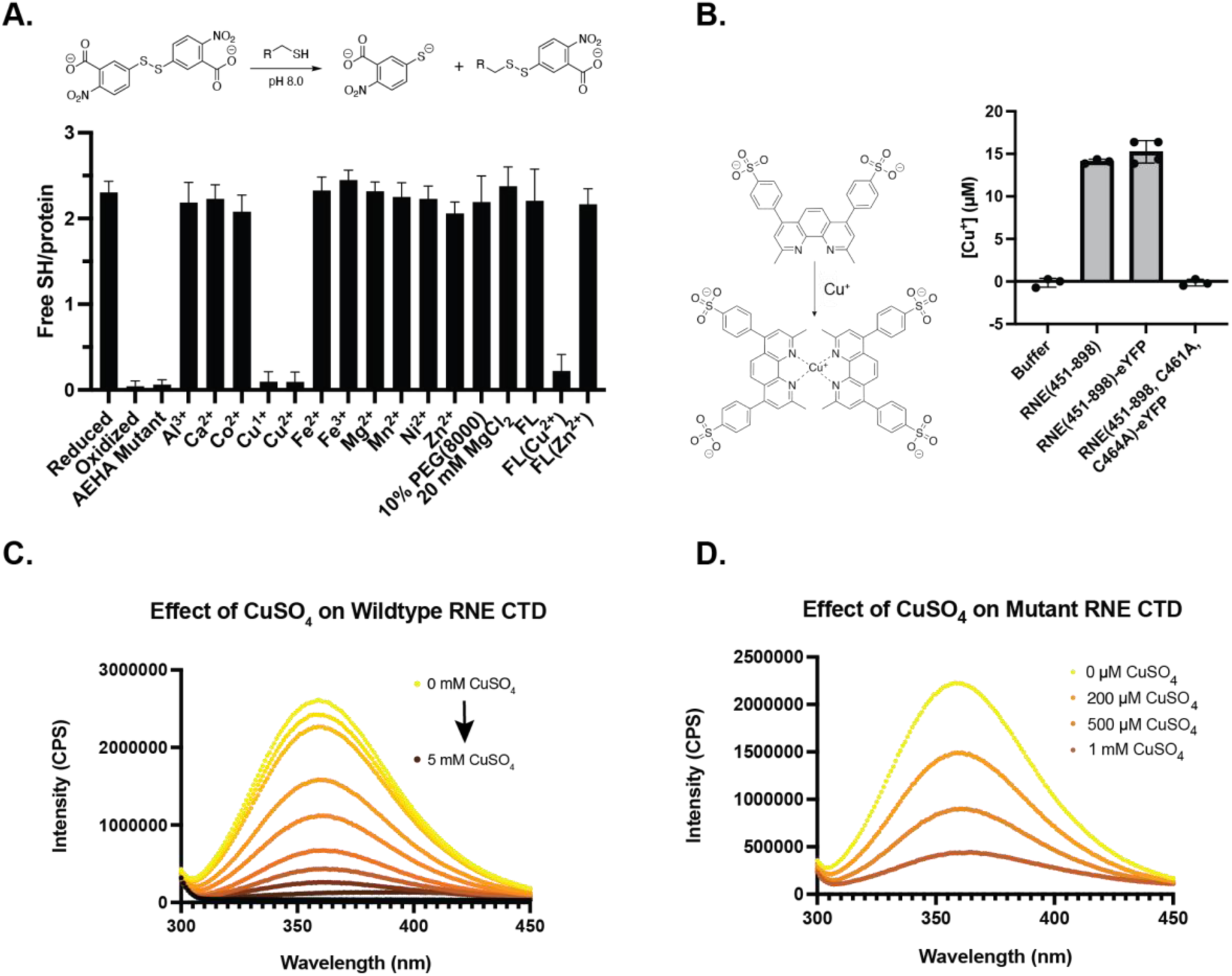
Copper interacts with RNase E through two distinct mechanisms: through cysteine-mediated reductive chelation of Cu(II) and a separate cysteine-independent binding mode. (A) An Ellman assay reveals a copper-specific interaction that limits C461 and C464 free thiol access. RNase E does not bind zinc in vitro but free thiol access is sensitive to disulfide bond formation. (B) Schematic representation of bathocuproine sulfonate (BCS) coordinating with Cu^1+^ in a BCS assay. Quantification of Cu^1+^ produced by reductive chelation of Cu^2+^, upon exposure to unlabeled RNase E (451-898) and RNase E(451-898)-eYFP, which consist of both cysteines versus RNase E(451-898, C461A, C464A)-eYFP in which both cysteines have been mutated to alanines. Quantification reveals that the concentration of Cu^1+^ substantially increases in the presence of both cysteines. (C) Unlabeled wildtype RNase E (451-898) titration by Cu^2+^ (Top to Bottom: 0 µM, 50 µM, 100 µM, 200 µM, 400 µM, 700 µM, 1 mM, 1.5 mM, and 5 mM CuSO_4_) monitored by protein intrinsic fluorescence intensity quenching with plot of fluorescence intensity vs. CuSO_4_ concentration. Increasing concentrations of CuSO_4_ result in a decrease in tryptophan fluorescence intensity. (D) Unlabeled RNase E (451-898, C461A, C464A) variant titration by Cu^2+^ monitored by protein intrinsic fluorescence intensity quenching. Increasing concentrations of CuSO_4_ result in a decrease in tryptophan fluorescence intensity.

We hypothesize that Cu^1+^ and Cu^2+^ may directly bind to RNase E or facilitate cysteine oxidation. If Cu^2+^ coordinates with cysteines via reductive chelation (Figure S4E), it should generate Cu^1+^, which can be quantified using the colorimetric chelator bathocuproine sulfonate (BCS). BCS binds Cu^1+^ and absorbs strongly at 483 nm. In addition, BCS does not interact with Cu^2+^, allowing for selective measurement of Cu^1+^. A standard curve using BCS and Cu^1+^ under reducing conditions was constructed to quantify the amount of Cu^1+^ (Figure S4F).

To determine whether RNase E reduces Cu²⁺ to Cu¹⁺, we desalted freshly reduced *Cc*RNase E into a buffer lacking reducing agents and incubated 10 µM RNase E with BCS following a 2-minute incubation with a 10-fold excess of Cu²⁺. This reaction yielded 15.2 ± 1.5 µM Cu¹⁺, whereas the control reaction without protein showed no detectable Cu¹⁺ formation (0.3 ± 0.4 µM) (Figure 4B). To confirm the role of cysteines in Cu²⁺ reduction, we tested RNase E (451-898, C461A, C464A)-eYFP under the same conditions. This variant produced no appreciable Cu¹⁺ (−0.1 ± 0.9 µM) (Figure 4B), suggesting that Cu²⁺ induces disulfide bond formation in RNase E. Together with the Ellman assay data, these results indicate that Cu²⁺ binding to RNase E leads to its reduction to Cu¹⁺, concomitant with RNase E disulfide bond formation.

### Zinc and copper do not mediate tetramerization of CcRNase E

Building on the Ellman and BCS assay results that revealed Cu^2+^ induces disulfide bond formation in RNase E (Figure 4A, 4B), we next explored whether there disulfide bonds were intermolecular leading to increases in oligomerization state. We analyzed RNase E (451-898) and RNase E (1-898, D403C) incubated with varying amounts of Cu^2+^, using native PAGE (Figure S4C, S4D) and RNase E (451-898) with non-reducing SDS-PAGE (Figure S4G). If intermolecular disulfide bonds were formed, we would expect tetramers under native conditions. In contrast, tetrameric (Cys)_4_Cu^1+^ complexes would maintain tetramerization in both assays, while Cu^1+^ complexes with two cysteines as (Cys)_2_Cu^1+^ would show no change in either assay. However, no appreciable change in the oligomerization state was detected under native conditions, indicating that intermolecular disulfide bonds were absent. Thus, the observed reduction in free thiol content is likely due to a combination of intramolecular disulfide bond formation and Cu¹⁺ complexes formed during reductive chelation.

### Cu^2+^ can interact directly with RNase E and does not require C461 or C464

Experiments at this stage suggest that Cu^2+^ binds to RNase E and can facilitate disulfide bond formation by reductive chelation. This leads to the key question of whether Cu^2+^ can coordinate cysteines or other nearby metal-binding residues. We, therefore, examined the quenching of the intrinsic tryptophan fluorescence intensity of the protein as a function of increasing concentrations of CuSO_4_. 50 µM unlabeled RNase E was incubated in a buffer of 100 mM NaCl and 20 mM Tris-Cl, pH 7.5, with and without varying concentrations of CuSO_4_. Decreasing fluorescence intensity was correlated with increasing concentrations of CuSO_4_ for unlabeled wild-type RNase E (451-898) (Figure 4C, S4A) and the unlabeled C461A/C464A variant (Figure 4D, S4B). This data suggests Cu^2+^ can interact with RNase E at alternative residues other than C461 or C464. Notably, these assays were done within a 20 mM Tris buffer that may compete with the protein for binding to copper. However, assays performed in the MOPS buffer confirmed a similar decrease in tryptophan fluorescence intensity, suggesting Tris does not interfere with RNase E-Cu(II) interaction (Figure S5A, S5B). These tryptophan fluorescence assays suggest that Cu(II) binds close enough to tryptophan to facilitate quenching or Cu(II) induces conformational changes in RNase E, altering the local chemical environment around tryptophan residues and reducing fluorescence intensity. Moreover, these assays suggest Cu(II) binds to RNase E in a C461/C464-independent manner.

To validate this tryptophan fluorescence quenching observations, we can directly investigate the interaction sites of Cu^2+^ using electron paramagnetic resonance (EPR) spectroscopy. The line shape of continuous wave (CW) EPR provides information on the identity of the atoms that are directed coordinated to Cu^2+^[30]. Notably, the spectrum of the RNase E wild-type and C461A/C464A mutant show almost identical peak positions (Figure 5A), indicating very similar Cu^2+^ binding in both protein samples. Note, that the line shape is distinct from that of free CuSO_4_ in buffer, showing that all Cu^2+^ is bound to RNase E. Interestingly, of the 700 µM Cu^2+^ added to the sample, only 155 µM Cu^2+^ for RNase E wild-type, and 306 µM Cu^2+^ for RNase E C461A/C464A mutant can be accounted for by EPR, suggesting that reductive chelation may generate (Cys)_2_Cu^1+^ species that are not visualizable by EPR.

**Figure 5.**
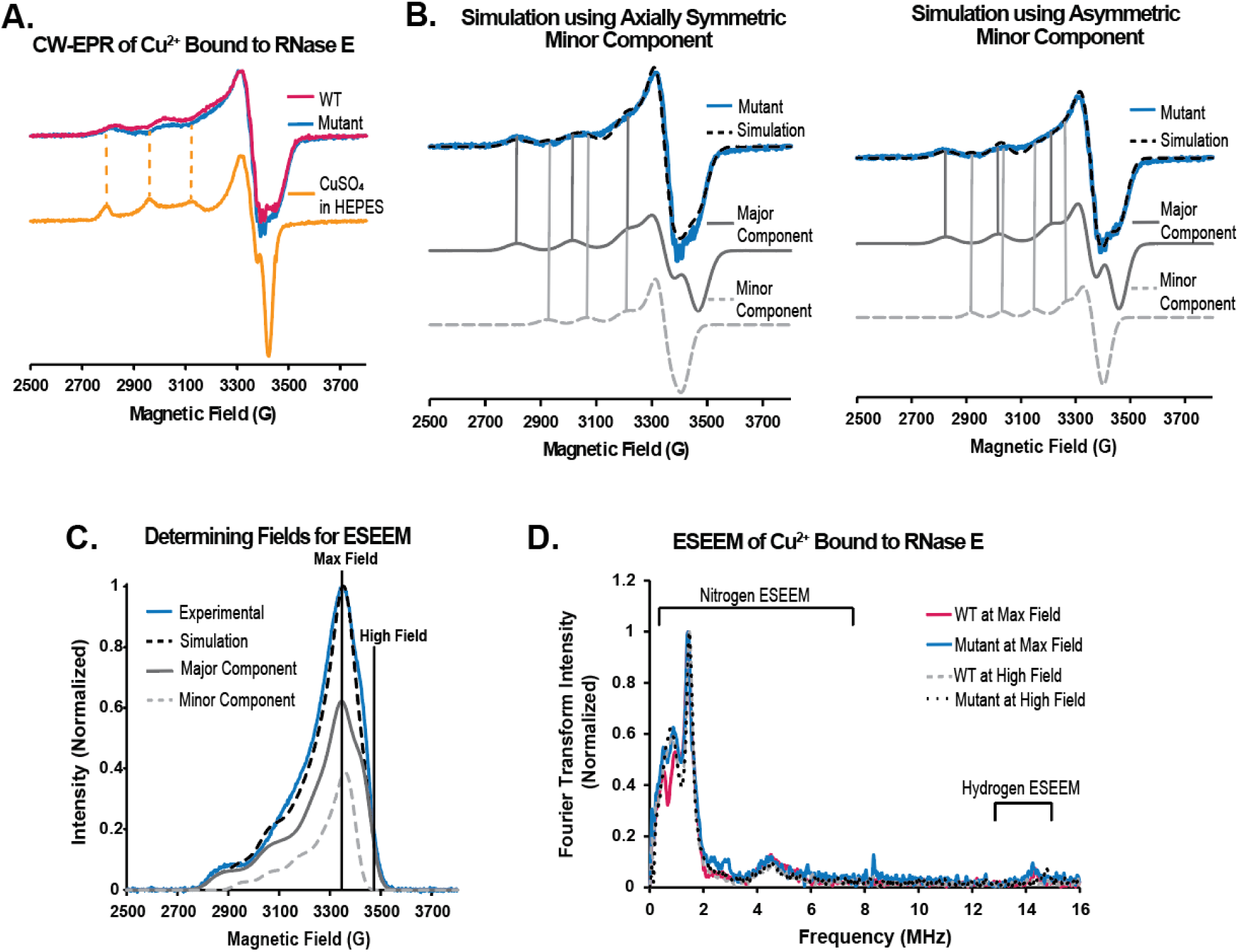
EPR provides support that RNase E binds to Cu^2+^ independent of cysteines and in association with the imidazole ring of a histidine. (A) CW-EPR spectra, collected at 80 K, of RNase E (451-898) wild-type (pink), the RNase E(451-898, C461A/C464A) variant (blue), and CuSO_4_ in HEPES buffer, pH 7.5 (yellow). The dashed yellow lines highlight the absence of peaks due to Cu^2+^ coordination to HEPES buffer in the protein samples. As the peak positions and intensities are almost identical in both RNase E samples, only the C461A/C464A variant data is shown in (B). CW-EPR spectra of RNase E wild-type data can be found in the SI. (B) Simulation of the C461A/C464A variant (blue) is overlaid in dashed black. The RNase E spectrum shows two components. The major component (grey) accounts for 70% of the simulation (dashed black), and the minor component (dashed light grey) accounts for the remaining 30% of the simulation (dashed black). The minor component was simulated using both axially symmetric (left) and asymmetric (right) coordination geometry. (C) Field swept spectrum of the RNase E C461A/C464A mutant (blue) overlaid with the integrated CW-EPR simulation (dashed black) using an asymmetric coordination geometry for the minor component (dashed light grey). ESEEM experiments were carried out at the field with the maximum intensity (3340 G) and a higher field (3463 G), highlighted with black lines. At the high field, only the major component (grey) contributes to the spectrum. (D) ESEEM spectra of RNase E wild-type and C461A, C464A mutant collected at the maximum intensity (pink and blue, respectively), RNase E wild-type collected at the high field (dashed grey), and RNase E mutant collected at the high field (dotted black). All spectra show peaks indicative of Cu^2+^ interacting with the imidazole ring of histidine.

Simulation of the data required two components (Figure 5B, S6A, S6B), which indicates two distinct Cu^2+^ binding sites on the protein. The major component (75% for wild-type and 70% for the mutant) in both samples indicate an octahedral, square planar, or square pyramidal metal binding site [30–33]. The major components in both proteins had a g_‖_ of 2.218 and A_‖_ of 187 G. By comparing these values to previously reported values for different combinations of nitrogen, oxygen, and sulfur coordination, it becomes clear that the major component is due to nitrogen and oxygen coordination in the equatorial plane [30, 33, 34]. Additionally, because the major component has the exact same g_‖_ and A_‖_ values in the wild-type and C461A/C464A variant, cysteines are not involved in the Cu^2+^ coordination of the major component.

The minor component of the RNase E wild-type and C461A/C464A variant (25% and 30%, respectively) also exhibited the same g-and hyperfine parameters. However, a distinct coordination geometry was less clear. When simulating using axially symmetric tensors, we found a g_‖_ of 2.203 and A_‖_ of 136 G (Figure 5B, left). These values do not align with expected values for an axially symmetric geometry [30] and A_‖_ values smaller than 140 G can point towards a trigonal planar or tetrahedral coordination [35–37]. Therefore, we also simulated the data using coordination for the minor component. This fit achieved a similar quality of fit to the data as the one that assumes an octahedral coordination (Figure 5B, right).

To further probe the binding sites of Cu^2+^, we performed electron spin-echo envelope modulation (ESEEM) experiments. ESEEM can observe the interaction between the electron spin of Cu^2+^ and the nuclear spin of elements within 4-8 Å of Cu^2+^[38, 39]. As the simulation of the CW-EPR data identifies two components, we performed the ESEEM experiments at two different fields[40]. The first field is at the maximum intensity of the field swept spectrum and samples both the major and minor components (Figure 5C). The second higher field samples only the major component. The resulting ESEEM spectra are shown (Figure 5D). The field swept spectrum for the RNase E wild-type and the raw time domain ESEEM spectra can be found in the SI (Figure S6C, S6D). Each spectrum shows three peaks below 2 MHz and one broad peak at 4 MHz, which indicates coordination to the imidazole ring of histidine[40–43]. Moreover, the integrated intensity of the peaks below 11 MHz compared to the integrated intensity of the peak at 14 MHz can be used to estimate the number of histidines that are coordinated to Cu^2+^[39, 40, 42–44]. Surprisingly, when measured at the maximum field, both the RNase E wild-type and C461A/C464A variant show coordination to one histidine. The spectra for both samples measured at the high field also shows coordination to only one histidine. The CW-EPR and ESEEM data confirm that in the major component, Cu^2+^ is directly bound to one histidine and no cysteine. Interestingly, the data also indicates that the minor component is likely also bound to one histidine residue.

Our data collectively indicate that Cu^2+^ can interact with RNase E in two distinct ways: cysteines can reductively chelate Cu^2+^ to form disulfide bonds, sequestering Cu as (Cys)_2_Cu^1+^ complexes, and Cu^2+^ can coordinate His residues directly.

### Copper solidifies RNase E condensates in a cysteine-dependent manner in vitro

Given the significant role of RNase E’s cysteines in phase separation in vivo (Figure 3), we investigated how these residues influence phase separation in vitro. We first assessed the effect of reducing conditions on the ability of RNase E to phase separate in the presence of MgCl_2_. To do so, we purified unlabeled wild-type CcRNase E (451- 898) and conducted phase separation assays by mixing 20 µM protein to produce conditions of 70 mM NaCl and 20 mM Tris with either 1 mM DTT only, 20 mM MgCl_2_ only, or both 1 mM DTT and 20 mM MgCl_2_. We also evaluated the phase separation behavior of the RNase E (451-898, C461A/C464A) variant under the same conditions (Figure 6A, *top*). Phase contrast microscopy revealed that wild-type CcRNase E (451-898) forms phase-separated droplets only in the presence of MgCl_2_ when reducing conditions are maintained. To optimize reducing conditions, a dose-response analysis in the presence of 20 mM MgCl_2_ revealed that concentrations of at least 1 mM DTT were necessary for robust phase separation of CcRNase E (451-898) (Figure 6B).

**Figure 6:**
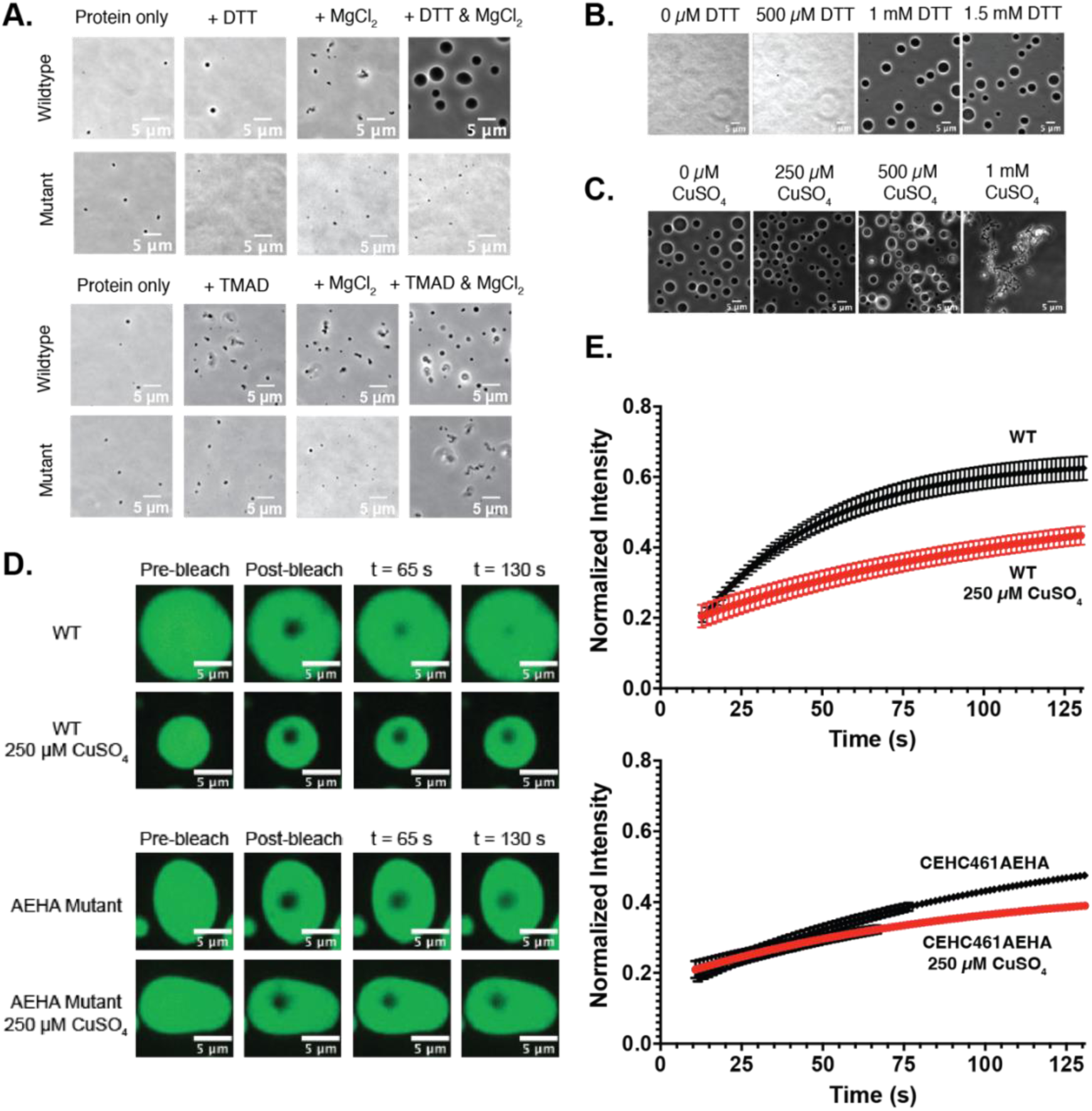
In vitro RNase E condensate material properties are regulated by copper binding and reducing conditions in a manner that is dependent upon C461 and C464. (A) Representative phase contrast microscopy imaging of unlabeled RNase E (451-898) reveals that RNase E phase separation with MgCl_2_ requires the presence of a reducing agent (DTT), while the presence of an oxidizing agent (TMAD) promotes aggregation. (B) Representative phase contrast microscopy images of unlabeled RNase E (451-898) demonstrate that an increase in DTT concentration positively correlates with an increase in RNase E phase separation. (C) Representative phase contrast microscopy images of unlabeled RNase E (451-898) further demonstrate that high concentrations of CuSO_4_ promote RNase E aggregation. The in vitro droplet imaging assays consist of 20 µM RNase E incubated in 20 mM Tris buffer, pH 7.4 and 70 mM NaCl in the absence or presence of varying concentrations of DTT and CuSO_4_, 1 mM TMAD, and 20 mM MgCl_2_. (D) and (E) The presence of CuSO_4_ impacts RNase E material properties, causing RNase E droplets in vitro to become more solid-like, resulting in slower recovery from fluorescence recovery after photobleaching (FRAP) for both wildtype and the C461A/C464A variant.

In the absence of the reducing agent, freshly desalted RNase E (451-898), stored in 20 mM Tris, pH 7.5, and 200 mM NaCl, forms only small aggregates, even in the presence of MgCl_2_ (Figure 6A, *top*). In contrast, the C461A/C464A variant did not form micron-scale-sized assemblies under any surveyed conditions, indicating the importance of cysteines and reducing conditions in the phase separation of RNase E. We further explored the effects of oxidizing conditions using 1 mM TMAD. Under these conditions, RNase E (451-898) formed smaller, dynamically arrested assemblies in the presence of MgCl_2_, suggesting that free cysteines are essential to the initiation and coarsening of RNase E assemblies (Figure 6A, *bottom*).

Building on previous in vivo observations, where CuSO_4_ caused RNase E foci in *C. crescentus* cells to diffuse at high concentrations, we examined its impact on RNase E phase separation in vitro. We titrated increasing amounts of CuSO_4_ (0 to 1 mM) into pre-formed RNase E droplets to assess changes in droplet morphology (Figure 6C). Imaging revealed that at concentrations above 500 µM, CuSO_4_ induced a dynamic arrest of RNase E droplets, resulting in aggregate-like assemblies.

To further characterize the effect of CuSO_4_ on the material properties of RNase E droplets, we measured fluorescence recovery after photobleaching (FRAP) of RNase E (451-898)-YFP assemblies. We examined whether CuSO_4_ causes wild-type RNase E (451-898)-YFP and the C461A/C464A variant to adopt a more solid-like state, as the aggregates observed at high CuSO_4_ concentrations suggested. FRAP analysis was conducted at 250 µM CuSO_4_ (Figure 6D). Increasing CuSO₄ concentrations resulted in slowed FRAP, suggesting a transition to more solid-like RNase E assemblies (Figure 6E).

We next considered whether this copper-stimulated change in RNase E material properties depended on C461 and C464. Unlike the wild-type RNase E protein, adding Cu^2+^ to RNase E (451-898, C461A/C464A)-YFP led to milder differences in the FRAP experiment. Moreover, using phase microscopy, we observed that wild-type RNase E (451-898), when combined with PNPase and poly(A) substrate, formed biomolecular condensates with diameters that ranged from 1.8 µm - 11.7 µm (n = 226) (Figure S7C). In comparison, in the presence of PNPase and poly(A), the RNase E (451-898) C461A/C464A variant only formed small assemblies ranging from 0.5-1.8 µm in diameter. This supports a model in which Cu^2+^ interactions at a site distal from the C461-C464 cysteine motif may facilitate networking interactions separate from disulfide bond formation, resulting in more solid-like RNase E condensate assemblies.

### RNase E condensates maintain PNPase activity in the presence of CuSO_4_

At this stage, we’ve observed that copper associates with RNase E condensate and can lead to more solid-like assemblies. Does this change in RNase E material properties arrest or even protect client protein function? To test the impact of copper associated condensates, we examined whether RNase E (451-898) protects PNPase activity from Cu²⁺ mismetallation. To do this, we utilized Thioflavin T, which exhibits increased Stokes shift and emission intensity in the presence of poly(A) [61]. Since the emission intensity scales with poly(A) concentration, it serves as a readout of PNPase activity. Without RNase E, PNPase degraded 77.5 ± 2.3% of poly(A) over 30 seconds when no Cu²⁺ was present. The addition of 125 µM CuSO₄ reduced degradation to 36.4 ± 5.6%, and 250 µM CuSO₄ further decreased it to 26.6 ± 1.2%. However, when RNase E was present, phase separation enhanced poly(A) degradation to 96.6 ± 9.4%, 94.9 ± 19.9%, and 43.3 ± 4.8% in the presence of 0 µM, 125 µM, and 250 µM CuSO₄, respectively (Figure 7A, S7D). These results suggest that at 125 µM, CuSO₄, RNase E condensates preserve the activity of PNPase. Whereas higher dosages of copper lead to more solid-like assemblies that also arrest PNPase function.

**Figure 7:**
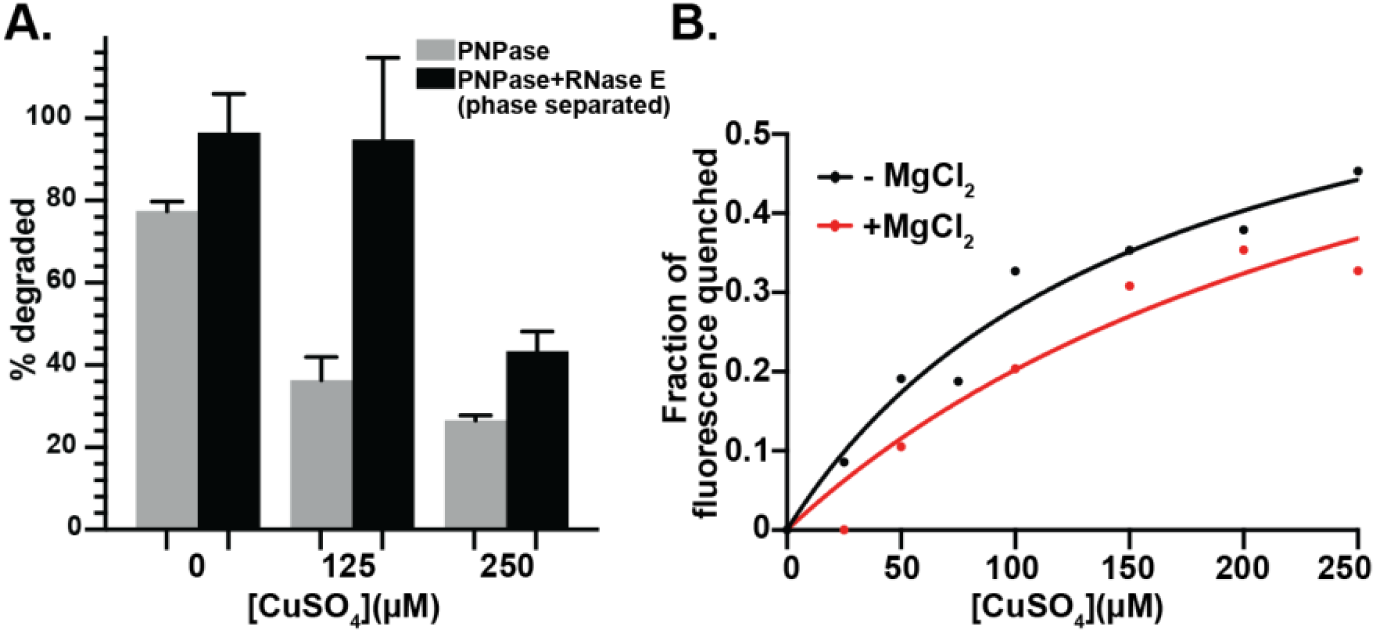
RNase E condensates preserve the ribonuclease activity of PNPase when exposed to CuSO4 in vitro. (A) Percent poly(A) degraded in the presence of PNPase with or without wild-type RNase E (451-898) using a fluorescent thioflavin-T tracking endpoint assay. RNase E progressively shields PNPase from Cu^2+^ mismetallation as Cu^2+^ concentration increases. (B) The intrinsic tryptophan fluorescence of 50 µM unlabeled PNPase in the presence or absence of 20 mM MgCl_2_ was monitored with increasing concentrations of CuSO_4_. The addition of the magnesium cofactor attenuates the Cu^2+^ dose-dependent reduction in PNPase’s intrinsic tryptophan fluorescence.

Based on the PNPase activity assays, we speculated that RNase E condensates may provide a protective zone for client proteins to perform biochemistry away from high copper stress. Indeed, copper toxicity in microbes often stems from mismetallation, where copper ions replace essential metal cofactors in proteins, impairing enzymatic functions. To investigate this, we examined copper binding to PNPase by monitoring changes in its intrinsic tryptophan fluorescence in the presence and absence of PNPase’s magnesium cofactor. With increasing concentrations of CuSO_4_, we observed a decrease in tryptophan fluorescence signal with an apparent Kd of 158.0 µM in the absence of magnesium and an apparent Kd of 301.9 µM in the presence of magnesium. Interestingly, the addition of magnesium attenuates the binding of CuSO_4_ to PNPase. This observation suggests that copper binds to PNPase in a manner that may be competitive with PNPase’s essential cofactor magnesium (Figure 7B, S7A, S7B).

## Discussion

Our findings support a model in which BR-bodies function as a molecular rheostat, dynamically adjusting their functional role to protect client proteins across varying copper levels (Figure 8). At low copper concentrations (<50 µM), RNase E condensates remain highly dynamic, supporting high cellular fitness when trace metals are depleted by chelators (Figure 2, S2B). Here’s a clearer version with two separate sentences: In a moderate copper range (100–250 µM), RNase E phase separation enhances cellular fitness (Figure 2F, S3B). In vitro, copper-induced condensates become more solid-like (Figure 6D) while preserving PNPase activity, which would otherwise be lost due to copper exposure (Figure 7). These findings suggest that BR-bodies create a protective microenvironment that shields PNPase and other client proteins from copper-induced inactivation, likely by providing competitive copper-binding sites that prevent mismetallation. However, at high copper concentrations (>500 µM), the condensate initially dissolves upon short exposure (Figure 2) before transitioning into stable, solid-like assemblies via copper-induced disulfide bond formation (Figures 4, S2C, 6). Under these conditions, cell viability is severely compromised, as seen in efficiency-of-plating assays (Figure S3B).

**Figure 8:**
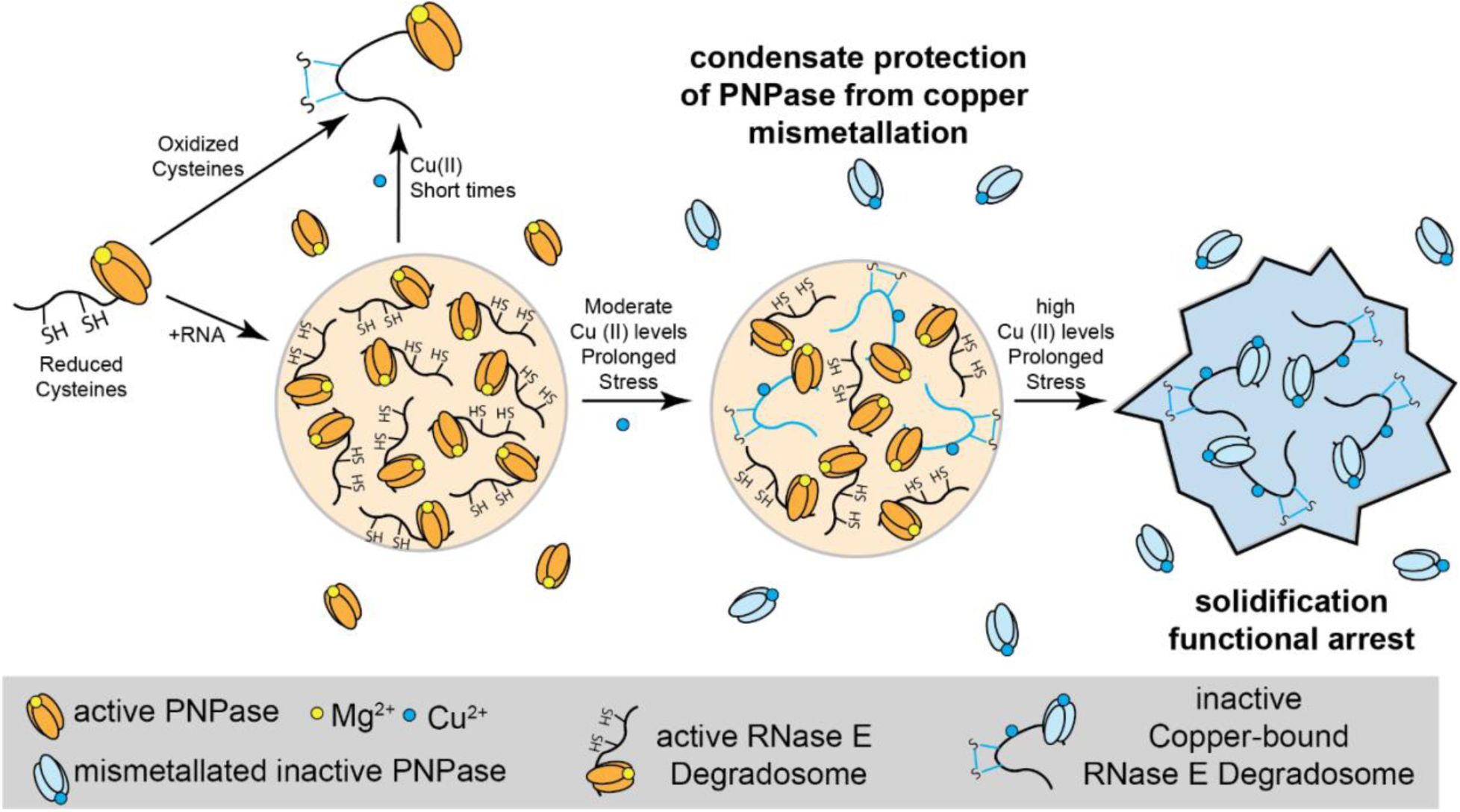
BR-bodies act as a molecular rheostat, dynamically responding to fluctuating metal stress. Under reducing conditions with cysteines and RNA, BR-bodies form robust, liquid-like condensates. Oxidation of cysteines by TMAD or copper drives RNase E into the dilute phase. At moderate copper levels (100–250 µM Cu²⁺), BR-bodies create protective microenvironments that shield PNPase from copper-induced inhibition. However, at high copper concentrations (>500 µM Cu²⁺), BR-bodies initially dissolve before solidifying over time through copper-induced RNase E disulfide bond formation, arresting their function. These adaptive responses enhance *Caulobacter* fitness under copper stress.

BR-bodies enhance cell fitness and preserve RNA decay activity in *Caulobacter*, potentially representing a broader design principle of how biomolecular condensates regulate metal-enzyme interactions under heavy metal stress. Indeed, a wide-variety of neurodegenerative disease associated condensates are mediated by metal ions [62–64]. The impact of condensates depends on the number of metal-binding sites, their affinity, and their saturation level. When binding sites are only partially saturated and have moderate affinity, condensates may create protective zones that prevent inhibitory metal binding to client proteins. Conversely, full saturation of weak binding sites can lead to high local metal ion concentrations, benefiting enzymes that require these metals but also increasing toxicity and mismetallation risks. As seen with RNase E, metal concentrations can dynamically alter the material properties of condensates (Figure 6D), influencing enzyme and substrate dynamics. This adaptable phase behavior allows condensates to sequester, protect, or modulate enzyme activity in response to environmental fluctuations. Moreover, caution should be considered with biochemistry studies of condensates, metals and use of the hexahistidine purification tag as this can drive synthetic condensate formation [65]. Collectively, understanding and applying these mechanisms could offer new insights into bacterial homeostasis and inform strategies for engineering stress-resistant cells in biotechnology and medicine.

## Methods

### Caulobacter crescentus cell growth and stress treatment

*Caulobacter crescentus* strains (*rne::rne-eyfp* or *parB::CFP-ParB; popZ::mCherry-popZ* or *rne::rne(C461A/C464C)-eyfp*) were streaked and incubated on a PYE agar plate supplemented with relevant antibiotics at 28 °C – 30 °C. When appropriate, the media was supplemented with either vanillate (5 μM), xylose (0.2%), gentamycin (0.5 μg/ml), kanamycin (5 μg/ml), chloramphenicol (2 μg/ml), spectinomycin (25 μg/ml), and/or streptomycin (5 μg/ml).

An overnight liquid PYE culture supplemented with the appropriate antibiotics was inoculated with an individual colony for copper stress tests. The culture was grown at 220 RPM at 28 °C to stationary phase at OD_600_ 0.9 – 1.0, measured at 600 nm in a cuvette using a UV-1600PC spectrophotometer (VWR). The culture was then split into equal volumes and supplemented with 500 µM CuSO_4_, 500 µM H_2_O_2_, 500 µM TMAD, 1 mM ZnCl_2,_ and 20 mM MgCl_2_. For the CuSO_4_ titration, the culture was split into equal volumes and diluted to OD_600_ 0.2-0.4 for imaging and supplemented with various CuSO_4_(ThermoFisher Scientific) concentrations: 0,100, 250, 300, 350, 400, 500, 700, 800, and 1000 µM. Cell stress treatments were performed by diluting the culture to OD_600_ 0.2-0.4 for imaging before supplementation with the desired copper stress. Cells were then immediately spotted on a 1.0% PYE agarose pad. Samples were imaged at short intervals between 4 and 8 minutes after adding copper stress. Imaging pads were supplemented with the same concentration of CuSO_4_ as was introduced to the aliquoted liquid culture. For the CuSO_4_ washout experiments, after CuSO_4_ stress imaging, the culture was spun down for two minutes; the supernatant was removed and washed 5 times over 30 minutes with fresh PYE media. Then, the culture was repleted with fresh media and imaged to monitor potential recovery from stress. Replacement strains containing a xylose-inducible copy of RNase E and a vanillate-inducible test construct were initially grown in media containing xylose overnight, then washed three times with 1mL growth media, and resuspended in growth media containing vanillate, diluted, and grown overnight. Stationary phase cultures were then diluted to 0.2-0.4 OD and spotted on PYE 1.5% agarose pads for imaging.

For the *Caulobacter* long-term copper exposure experiments, *C. crescentus* were diluted 1:100 from an overnight modified HIGG (mHIGG, see below) culture supplemented with appropriate antibiotics. Cells were grown to OD_600_ 0.2-0.3 at 28 °C. For strain JS49, RNase E (1-898)-eYFP is under the control of the *vanA* promoter, and cells were induced with 0.5 mM vanillate (pH 8.0) for 1 hr. Cells were immobilized on a 1.0% mHIGG-agarose pad before imaging. To induce moderate copper stress at long times, strains were incubated with 1.6 mM CuSO4 for 1 h after vanillate induction at 30 °C. Cells were then immobilized on a 1.0% M2G-agarose or 1.0% mHIGG-agarose pad supplemented with 1.6 mM CuSO_4_. HIGG media was modified to generate mHIGG: 9.0 mM L-arginine pH 7.0, 9.0 mM NH_4_Cl pH 7.0, 8.32 mM glucose, 5 mM imidazole pH 7.0, 1.45 mM CaCl_2_, 610.4 µM KH_2_PO_4_, 389.6 µM Na_2_HPO_4_, 240 mM MgCl_2_, 38.2 µM ZnSO_4_, 25.1 µM FeSO_4_, 10.5 µM nitrilotriacetic acid, 9.12 µM MnSO_4_, 6.72 µM Na_2_EDTA, 1.57 µM CuSO_4_, 852 nM Co(NO_3_)_2_, 464 nM B(OH)_3_, 150 pM (NH_4_)_6_Mo_7_O_24_.

### E. coli cell growth

*Escherichia coli* BL21(DE3) or TOP10 strains were grown at 37 °C and cultured in an LB medium (Sigma), supplemented with kanamycin (30 µg/mL) or ampicillin (50 µg/mL). For induction, cells were induced with isopropyl-β-D-thiogalactopyranoside (IPTG) (1 mM) for two hours or 0.0004% arabinose for one hour. Strains were analyzed at mid-exponential growth phase (OD_600_ 0.3-0.6), using a UV-1600PC spectrophotometer (VWR).

### Fluorescence microscopy and phase imaging

For cell imaging, bacterial cells were immobilized on 1.0% agarose pads made with PYE or LB media on microscope slides (3051, Thermo Scientific). Images were obtained utilizing a Nikon Eclipse Ti-E inverted microscope equipped with an Andor Ixon Ultra DU897 EMCCD camera and a Nikon CFI Plan-Apochromat 100×/1.45 Oil objective and intermediate 1.5× magnification. Carl Zeiss Immersol 518 oil was used. The excitation source was a Lumencor SpectraX light engine. The filter sets utilized for YFP, CFP, and mCherry imaging were chroma 96363, 96361, and 96322 models. YFP was exited at 512 nm, mCherry was excited at 549 nm, and emission was monitored using the CFP/YFP/mCherry Chroma filter cube. Images were acquired with Nikon NIS-Elements AR software.

For in vitro droplet imaging assays, 20 µM RNase E was incubated in 20 mM Tris buffer, pH 7.4 and 70 mM NaCl in the absence or presence of varying concentrations of DTT and CuSO_4_, 1 mM TMAD, and 20 mM MgCl_2_.Imaging samples were pipetted into a 1 mm well formed by an adhesive spacer (Electron Microscopy Sciences) affixed to a microscope slide (VWR) and sealed with a glass coverslip (VWR). Slides were inverted and allowed to sit at room temperature for 30 min before on a Nikon Eclipse Ti-E inverted microscope with a Plan Apo-(lambda) 100×/1.45 oil objective and 518F immersion oil (Zeiss). Images were taken with an Andor Ixon Ultra 897 EMCCD camera.

#### PNPase Degradation Assay by Microscopy

Fluorescence microscopy samples were prepared by thawing requisite proteins on ice and mixing a buffer to create a final concentration of 20 mM Tris pH 7.5, 20 mM MgCl_2_, 70 mM NaCl, 0.5 mM DTT, 5 μM purified PNPase, 25 ng/μL poly(A) RNA and 20 μM purified wildtype RNase E (451-898) or the C461A/C464A RNase E (451-898) variant. Imaging samples were pipetted into a 1 mm well formed by an adhesive spacer (Electron Microscopy Sciences) affixed to a microscope slide (VWR) and sealed with a glass coverslip (VWR). Slides were inverted, and samples lacking PNPase were allowed to sit at room temperature for 30 minutes before imaging. Samples with PNPase and poly(A) were also imaged within 30 min of plating on a Nikon Eclipse Ti-E inverted microscope with a Plan Apo-(lambda) 100×/1.45 oil objective and 518F immersion oil (Zeiss). Images were taken with an Andor Ixon Ultra 897 EMCCD camera.

### Fluorescence image processing

Cell image analysis was conducted using the MicrobeJ software [46]. A set of images of each unique condition was uploaded to MicrobeJ, consisting of a combination of phase contrast, YFP, CFP, and mCherry channels when appropriate. The phase image was utilized to create a cell outline using the Medial Axis setting. For automated foci detection, the maxima foci function of microbeJ was used, and the tolerance setting was manually adjusted between 250 and 1000 to identify and outline foci on a test image. The Z-score, a parameter indicative of the number of standard deviations from the mean, was set at 20.0. The segmentation tool was then utilized to split adjoined foci, and aberrant foci with an area < 0.01 µm^2^ and length > 1 µm were omitted. A Welch’s 2-tailed t-test using unequal variances was subsequently performed on all unique samples compared to the control. For each set of images uploaded, the same tolerance/Z-score parameters were used to quantify the average foci/cell. To determine partition ratios, we background-subtracted the RNase E maximum (foci) and minimum intensities within the cell bodies as calculated by microbeJ. We divided the maximum intensity of the YFP signal in all foci by the YFP intensity within the cell apart from the foci.

Additional details: Channel 1 was designated as “Bright” and the following parameters were selected: “Exclude on Edges”, “Shape descriptors”, “Segmentation”, “Edge Correction”, “Intensity” and “Shape”. Additionally, the Area was bound by 0.2-max µm^2^, Length 1-max, Width 0-max, Circularity 0-1 and Curvature, Sinuosity, Angularity, Solidity, and Intensity 0-max. Channel 2 was designated as “Dark” and the following parameters were selected: Area was bound by 0-max µm^2^, Length 0-2.81, Width 0-max, Circularity 0-0.74 and Intensity 1710-max.

### mRNA half-life measurements

*C. crescentus* NA1000 cells were grown in liquid PYE at 28°C to an optical density OD_600_ ∼ 0.3-0.6 (exponential phase). Copper II Sulfate was added to the cultures at the final concentration of 0 mM, 125 mM, or 500 mM for 8 minutes before the RNA extraction. Regarding *JS38* and *JS802*, cells were first grown overnight in liquid PYE + Kan + Gent + 0.2% xylose at 28 °C. Once the OD reached ∼0.3-0.6, cells were washed 3 times with PYE and resuspended in 25 mL PYE at an OD of 0.05. 500 μM Vanillate was added, and the cultures were incubated for 8 hours before the RNA extraction.

1 mL of cells was taken at time zero (before the addition of rifampicin) and vortexed with mL of RNAprotect Bacterial reagent. Following the addition of rifampicin (200 µg/mL), 1 mL aliquots were collected at predetermined intervals (1, 2, 4, and 8 minutes) and vortexed with 2 mL of RNAprotect Bacterial reagent as previously. Cells were then pelleted by centrifugation at 6000 rpm for 2 minutes and resuspended in 1 mL 65 °C pre-warmed TRIzol. The samples were then incubated at 65°C for 10 minutes. Afterward, 200 uL of chloroform was added to the samples, and the tubes were inverted 6-7 times. The samples were incubated for 5 minutes at room temperature before being centrifuged for 10 minutes at maximum speed. The temperature of TRIzol was set at 65 °C. After centrifuging the cells in RNAprotect for two minutes at 8000 rpm, they were resuspended in one milliliter of TRIzol that had been warmed up beforehand and incubated for ten minutes at 65 °C. The upper aqueous phase was then pipetted into a new tube. 2 uL of 1 mg/mL glycogen and 700 uL of isopropanol were added to the samples and incubated at −80 °C overnight. The following day, the samples were centrifuged for 1 hour at maximum speed at 4 °C. The supernatant was discarded, and the RNA pellets were washed with 1 mL of ice-cold 80% ethanol. The samples were then centrifuged for 10 minutes at maximum speed. The ethanol was discarded, and the RNA pellets were air-dried and resuspended in 50 uL of RNA resuspension buffer (10 mM Tris-HCl, pH 7.0, 0.1 mM EDTA).

For measuring mRNA half-lives by RT-qPCR, a master mix was prepared, including 1× Luna Universal One-Step Reaction Mix, 1× Luna WarmStart RT Enzyme Mix, 0.4 µM of forward and reverse primers, and Milli-Q water. 100 ng/uL of RNA from each collected sample was aliquoted into a 96-well plate and mixed with 19 uL of the master mix. The mixture was briefly vortexed and centrifuged. A QuantStudio Real-Time PCR machine was used to conduct the RT-qPCR reactions. RNA decay rates were determined by fitting a linear curve to the natural logarithm of the proportion of RNA that remained at each time point. Each time point’s RNA quantity was standardized to the 100% amount found in the time zero samples. The mRNA half-life was calculated using linear regression of the ln (% of RNA remaining) at each RNA extraction time point. The slope was then converted to a half-life measurement using the formula t1/2 = −ln(2)/slope.

#### Ellman Assay

Reaction buffer (100 mM Tris-Cl, pH 8.0) was prepared and used to generate stocks of DTT (1.5 mM) and 5,5’-dithiobis(2-nitrobenzoic acid) (DTNB, 5 mM in 50 vol.% DMSO). All reagents were stored on ice before use. A set of DTT standards were created covering 0 - 100 µM DTT were dissolved in reaction buffer before 200 µM DTNB was added and allowed to incubate at room temperature for 15 minutes. The absorbance at 410 nm (A_410_) was measured on a NanoDrop 2000C, and a line of best fit was generated: A_410_ = 0.001468·[SH]+0.0007433 (R2 = 0.9984), where [SH] represents the concentration of free sulfhydryl groups in µM. Error bars represent the results of two replicates.

For each condition, 5-10 µM RNase E (451-898), RNase E (451-898, C461A/C464A) [AEHA mutant], or RNase E (1-898, D403C) [FL-RNE] was incubated with 200 µM DTNB for 15 min in 100 mM Tris-Cl, pH 8.0 before A_410_ measurement. Protein was mixed with either a 5-fold excess of tris(2-carboxyethyl)phosphine (TCEP) for 30 min (reduced; AEHA mutant), a 20-fold excess of tetramethylazodicarboxamide (TMAD) for 1 hour (oxidized), a 10-fold excess of metal for 2 min, or phase separation was induced with either 10 wt% PEG (8000) or 20 mM MgCl_2_ before DTNB addition. Significance was determined using a one-way ANOVA compared to the results of RNase E (451-898).

Metal stocks were diluted immediately before use in 100 mM Tris-Cl, pH 8.0, except for CuCl, which was diluted into 7.5 M NH_4_OAc. Metal sources included AlCl_3_, CaCl_2_, CoCl_2_, CuCl, CuSO_4_, FeSO_4_, FeCl_3_, MgCl_2_, MnCl_2_, NiSO_4_, and ZnSO_4._ The concentration of sulfhydryl groups was divided by the concentration of RNase E to obtain the number of sulfhydryl groups per protein. An error was determined based on two replicates.

#### Bathocuproine sulfonate assay for Cu^1+^

Reaction buffer (100 mM Tris-Cl, pH 8.0, 1 mM DTT) was prepared and used to dilute the 10 mM bathocuproine sulfonate (BCS). CuCl stocks (100 µM – 100 mM) were generated by dilution into 7.5 M NH_4_OAc. All reagents were stored on ice before use. A set of Cu^1+^ standards was created in triplicate covering 0 – 100 µM Cu^1+^ dissolved in reaction buffer before 500 µM DTNB was added and allowed to incubate at room temperature for 2 minutes. The absorbance at 483 nm (A_483_) was measured on a NanoDrop 2000C, and a line of best fit was generated: A_483_ = 0.001326·[Cu^1+^] + 0.0009528 (R^2^ = 0.9983), where [Cu^1+^] represents the concentration of free Cu^1+^ in µM. Error bars represent the results of two replicates. To measure the Cu^1+^ content after incubation of Cu^2+^ with RNase E in solution, the protein was treated with a 5-fold excess TCEP before being desalted into 200 mM NaCl, 100 mM Tris-Cl, pH 8.0..Protein was then incubated with a 10-fold excess of CuSO_4_ in 200 mM NaCl, 100 mM TrisCl, pH 8.0 for 2 minutes before adding BCS. For each 10 µM RNase E sample, 500 µM BCS was added, and an A_483_ measurement was immediately taken. An error was determined based on two replicates.

#### Thioflavin T assay for tracking poly(A) degradation

For reactions without RNase E, samples were prepared in the following order, with final concentrations as listed: 20 mM Tris-Cl (pH 7.5), 70 mM NaCl, 2.5 mM PNPase, 500 µM MgCl₂, 25 ng/µL poly(A), 10% PEG-8000, and 4 mM Na_2_HPO_4_ (phosphate). Phosphate was added last to initiate the reaction, which proceeded for 30 seconds before quenching. The reaction was quenched with final concentrations of 100 mM EDTA, 25 µM Thioflavin T, and 556 mM NaCl.

RNase E was added before PNPase to a final concentration of 20 µM for reactions with phase separation. In reactions with copper, CuSO₄ was introduced after PEG-8000, reaching final concentrations of either 125 µM or 250 µM.

To generate a standard curve for each condition, MgCl_2_ was omitted, and intensity measurements were taken with the following poly(A) concentrations: 20 ng/µL, 10 ng/µL, 5 ng/µL, and 0 ng/µL.

Fluorescence measurements were conducted using a Tecan Infinite M1000 microplate reader (Tecan Group Ltd., Männedorf, Switzerland). Samples were prepared in a 96-well U-bottom microplate (Greiner Bio-One, Kremsmünster, Austria). Fluorescence detection was performed in top-read mode with an excitation wavelength of 438 nm and an emission wavelength of 491 nm, both with a bandwidth of 5 nm. The instrument was set to manual gain mode with a fixed gain of 135. Each well was measured using 50 flashes at 400 Hz, and the Z-position was set to 18,400 μm to optimize signal detection.

To calculate the percentage of poly(A) degraded, ten intensity measurements were taken for three replicates of each condition after the reactions were quenched. The mean intensities for each replicate were associated with the poly(A) concentrations according to the standard curve fits of each condition. The standard deviations of each intensity measurement were calculated by propagating the uncertainties through a first-order Taylor expansion method given by:

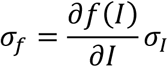

Here, *f*(*I*) the concentration of poly(A) based upon the standard curve fit *I* is the measured mean intensity from a particular replicate and *σ*_*I*_ is the standard deviation of *I*. Finally, given a starting poly(A) concentration of 25 ng/µL:

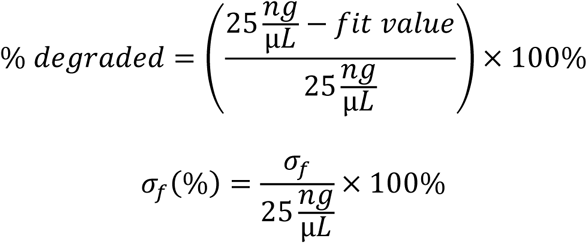

#### Fluorescence recovery after photobleaching (FRAP)

FRAP experiments were conducted using a Nikon Ti2 microscope equipped with a Yokogawa CSU spinning disk confocal system and an Andor DU-897 EMCCD camera. A Plan Apo λ 100× oil-immersion objective (NA 1.45) was used to acquire images with a pixel resolution of 512 × 512. The sample was illuminated using a LU-NV NIDAQ multi-laser system, with the 515 nm excitation laser line set at 40% power. The emission was collected through a 593.5 nm emission filter and a dichroic mirror (86006 - CFP/YFP/DsRed).

Images were captured at a 70 ms exposure time. The EMCCD camera was set to a gain multiplier of 80, a readout speed of 17 MHz, and a vertical shift speed of 3.3 µs with a vertical clock voltage amplitude of +4V.

For FRAP, a region of interest (ROI) within the condensates was photobleached using a Galvo XY scanning system. Fluorescence recovery was monitored over 63 time points.

The time interval between acquisitions was 10 seconds, and the acquisition rate was approximately 47.5 frames per second.

Data analysis was performed by normalizing fluorescence intensities to pre-bleach values and fitting the recovery kinetics using a single exponential model.

#### PNPase-mediated RNA-degradation assay

RNA degradation assays were performed at room temperature in 20 mM Tris–HCl (pH 7.5), 70 mM NaCl, 20 mM MgCl_2_, 4 mM Na_2_HPO_4_ (pH 7.5), and 0.5 mM DTT with 5 μM purified PNPase and 20 μM purified unlabeled wildtype RNase E CTD or the unlabeled C461A/C464A RNase E variant, in addition to 100 µM, 250 µM or 500 µM CuSO_4_ when appropriate. Reactions were initiated by adding 5 µM PNPase to a mixture containing 25 ng/μL poly(A) RNA.. For time-course assays, aliquots were withdrawn and quenched in 100 mM EDTA. Samples were denatured in 1.5 volumes of 2× RNA loading buffer containing 95% formamide, 18 mM EDTA, and 0.025% SDS and incubated at 95 °C for 3 min. Quenched ribonuclease reaction aliquots were loaded onto a pre-run 6% acrylamide gel containing 7 M urea in 1× TBE buffer (89 mM Tris base, 2 mM EDTA, 89 mM boric acid). To separate RNA, the gel was run in 1× TBE at 250 V at room temperature. Subsequently, the gel was rinsed in Milli-Q water for 5 min and stained for 20 min with 1× SYBR gold nucleic acid stain (Invitrogen) in 1×TBE. Each gel assay included RNA-only and protein-only controls. Gels were imaged using the BioRad ChemiDocTM MP imager with SYBR gold settings and quantified using the BioRad ImageLab software. The intensity of the protein-only lane was subtracted from each timepoint lane intensity and converted to ng of poly(A) in correlation with intensity based on the intensity in the lane with the starting amount of poly(A) and plotted against time (sec). The degradation rate was calculated using the initial time points up to 60 seconds for which a line of best fit was generated. From this equation, the slope was obtained to correlate to a degradation rate of poly(A) (ng/s). This rate was then converted to ng RNA/min/mg PNPase by multiplying this rate by 60 sec/min and dividing it by the amount of protein in mg.

#### Protein expression and purification of PNPase

Protein expression and purification were completed as described previously (Michael J. Collins et al., 2023). Briefly, plasmid pMJC0094 was constructed to heterologously express an N-terminally His_6_-tagged *C. crescentus* PNPase in *Escherichia coli*. Plasmid pMJC0094 was transformed into chemically competent Rosetta (DE3) cells, plated onto LB-Miller plates supplemented with 50 µg/mL ampicillin, and incubated overnight at 37 °C. An overnight 60 mL LB-Miller culture was inoculated from a single colony incubated at 37 °C. From this saturated culture, 6 L of LB-Miller media) was inoculated with 60 mL of the saturated culture and grown at 37 °C to mid-log phase (OD_600_ ∼0.5). The expression of PNPase was induced with 333 μM isopropyl-β-<u>D</u>-1-thiogalactopyranoside (IPTG) for 4 h at 25 °C. The cells were collected by centrifugation at 4 °C, 4000×*g*, for 30 min. The resulting pellet was washed with 60 mL resuspension buffer (50 mM Tris pH 7.5, 500 mM NaCl) before being pelleted at 4 °C, 4000×*g*, for 20 min and stored at - 80 °C. The cell pellet was thawed on ice and then resuspended in 10 mL lysis buffer per liter of culture (20 mM Tris HCl pH 7.5, 500 mM NaCl, 5 mM imidazole, 200 U benzonase) supplemented with SigmaFast™ protease inhibitor tablets (Sigma). The cell suspension was lysed by continuous passage through an Avestin Emulsiflex-C3 at 15,000 psi for 15 min at 4 °C. Cell debris was pelleted by centrifugation at 20,000×*g* for 45 min at 4 °C. The supernatant was loaded onto a HisTrap FF column (GE Healthcare) and washed with 20 column volumes of wash buffer (20 mM Tris HCl pH 7.5, 500 mM NaCl, 5 mM imidazole). Then, it was eluted with elution buffer (20 mM Tris HCl pH 7.5, 500 mM NaCl, 500 mM imidazole). The His_6_-tag was cleaved off with TEV protease. Fractions containing PNPase were supplemented with 50 mM sodium phosphate (pH 7.5) at 37 °C for 1 h to drive phosphorolysis of co-purifying RNA. The fractions were loaded onto a G-SepTM 6–600 kDa Size Exclusion Columns (G-Biosciences) and eluted with storage buffer (20 mM Tris HCl pH 7.5, 200 mM NaCl, 5% (v/v) glycerol). Fractions containing PNPase were concentrated using 50,000 MWCO Amicon centrifugal filters to 15.5 mg/mL (MW: 79530 g/mol, ε: 42985 1/M·cm), aliquoted, and flash-frozen in liquid nitrogen and stored at −80 °C.

#### Purification of RNase E-CTD

*C. crescentus* RNase E (451-898) was expressed in *E. coli* Rosetta BL21(DE3) from a plasmid (pET28::His6-MBP-TEV-RNase E (451-898)/ WSC1654) possessing a His_6_-tag and MBP fusion. Protein expression was induced in cells grown to mid-log phase (OD_600_ ∼0.5) at 37 °C with 1 mM IPTG for 4 hours before harvesting. Cells were frozen at −80 °C prior to purification after washing in 500 mM NaCl, 50 mM HEPES-KOH, pH 8.0, and re-pelleting. Harvested frozen cells were thawed on ice and then resuspended in 10 mL lysis buffer per liter of culture (20 mM Tris HCl pH 7.5, 500 mM NaCl, 5 mM imidazole, 1 mM β-mercaptoethanol, 5% glycerol, 100 U DNase I, 0.1% Triton X-100) supplemented with SigmaFast™ protease inhibitor tablets (Sigma). The cell suspension was lysed by continuous passage through an Avestin Emulsiflex-C3 at 15,000 psi for 45 min at 4 °C. Cell debris was pelleted by centrifugation at 20,000*g* for 45 min at 4 °C. The supernatant was loaded onto a HisTrap FF column (GE Healthcare) and washed with 20 column volumes of wash buffer (5 mM Tris HCl pH 7.5, 500 mM NaCl, 5 mM imidazole, 1 mM β-mercaptoethanol, 5% glycerin). Subsequently, the purification was eluted with elution buffer (20 mM Tris HCl pH 7.5, 200 mM NaCl, 200 mM imidazole, 1 mM beta-mercaptoethanol, 5% glycerin). The His_6_-tag and MBP-tag were cleaved off with TEV protease before being loaded onto a 16/60 Gel Filtration column to separate MBP from RNase E. Fractions containing RNase E were loaded onto a G-SepTM 6–600 kDa Size Exclusion Columns (G-Biosciences) and eluted with storage buffer (20 mM Tris-HCl, pH 7.5, 200 mM NaCl). RNase E’s fractions were concentrated using 30,000 MWCO Amicon centrifugal filters (MW: 49659 g/mol, ε: 32095 1/M·cm), aliquoted, and stored at - 80 °C.

#### RNase E-CTD EPR Sample Preparation

Purified RNase E protein was thawed, and buffer exchanged by a 2-hour followed by an overnight dialysis in HEPES buffer (20 mM HEPES pH 7.5, 200 mM NaCl). The protein stock was then concentrated to 267.04 µM. CuSO_4_ was added and allowed to incubate at room temperature for 30 minutes. CW-EPR samples for analysis were then prepared to a total volume of 100 µL with 20 vol% glycerol at a final RNase E concentration of 120 µM. Samples were then flash-frozen in liquid MAPP gas and stored at −80 °C.

#### EPR Experiments

Continuous Wave EPR: CW-EPR experiments were performed at 80 K with a Bruker E580 X-Band FT/CW spectrometer, a Bruker ER4118X-MD5 resonator, Oxford CF935 dynamic continuous-flow cryostat, and Oxford LLT 650 low loss transfer tube. The spectra were collected at a microwave frequency of ∼9.67 GHz with a 30 dB attenuation, 4 G modulation amplitude, 100 kHz modulation frequency, and 20.48 ms conversion time. The magnetic field sweep was centered at 3100 G and was 2000 G long, collected with 1024 data points and 200 scans. All simulations were performed using the pepper function in EasySpin Software [45].

Electron Spin-Echo Envelope Modulation EPR: Three-pulsed ESEEM experiments were performed at 18 K with a Bruker E580 X-Band FT/CW spectrometer, a Bruker ER4118X-MD4 resonator, Oxford CF935 dynamic continuous-flow cryostat, and Oxford LLT 650 low loss transfer tube. A pulse sequence of π/2 – τ - π/2 – T - π/2 – echo with a π/2 pulse width of 10 ns was used, where t = 140 ns, T = 280 ns, and was incremented in steps of 16 ns. The ESEEM experiments were carried out at 3340 G and 3480 G. The raw experimental data was background-subtracted using a stretched exponential decay, and the Fourier transformed time domain signal was calculated using the Bruker XEPR software.

#### Tryptophan Fluorescence Quenching Analysis

Tryptophan fluorescence data was obtained using a FluoroMax-3 fluorimeter (Jobin Yvon Horiba). Samples with unlabeled wild-type or mutant C461A/C464A RNase E (451-898) with or without PNPase were prepared by incubating 50 µM RNase E and 50 µM PNPase in 100 mM NaCl, 20 mM Tris, pH 7.4 with varying amounts of CuSO_4_ in a total volume of 70 µL. Measurements were taken using a quartz cuvette. The FluorEssence software was used to collect the data (Jobin Yvon Horiba), with an excitation wavelength set to 280 nm and emission wavelengths scanned between 300-450 nm. Maximum fluorescence intensities at 359, 358, and 350 nm were used to estimate the apparent K_D_ of Cu^2+^ interactions with RNase E (451-898), AEHA Mutant RNase E (451-898), and PNPase, respectively.

#### Efficiency of Plating (EOP) Assays

All *Caulobacter crescentus* strains used in this study were derived from the wild-type strain NA1000 (ΔCTD and ΔDBS as the only copy of RNase E on the chromosome which do not pick up suppressor mutations) and were grown at 28 °C in peptone-yeast extract (PYE) medium (Schrader and Shapiro, 2015). When appropriate, media was supplemented with vanillate (5 µM), xylose (0.2%), gentamycin (0.5 µg/mL), kanamycin (5 µg/mL), chloramphenicol (2 µg/mL), spectinomycin (25 µg/mL), and/or streptomycin (5 µg/mL) was added. Strains were analyzed at stationary (OD 0.9-1.2) and at exponential growth phase (OD 0.1-0.3). Optical density was measured at 600 nm in a cuvette using a UV-1600PC spectrophotometer. For the efficiency of plating (EOP) assays, cells were grown overnight in PYE, which lacked antibiotics and was diluted to OD_600_ 0.05. 10-fold serial dilutions were then performed and spotted on plates with a specified concentration of ethanol or CuSO_4_. After spotting cells, the plates were allowed to sit with the agar side down for 30 min to dry before the plates were incubated, inverted, at 28 °C for 1-3 days.

All *Agrobacterium tumefaciens* strains used in this study (JS5 [Al-husini mol cell 2018 ref] and JS376) were derived from the wild-type strain C58 and were grown at 28 °C in LB medium supplemented with gentamycin (30 µg/mL). Strains were analyzed at exponential growth phase (OD 0.3). For the efficiency of plating (EOP) assays, cells were grown overnight in LB + gent and were diluted to OD_600_ 0.05. 10-fold serial dilutions were then performed and spotted on LB plates either lacking CuSO_4_ or containing 3 mM CuSO_4_. After spotting cells, the plates were allowed to sit with the agar side down for 30 min to dry before the plates were incubated, inverted, at 28 °C for 1-3 days.

### JS801 rne::rneΔDBS SpecR

The CTD region lacking the degradosome binding sites (DBS) of RNase E (RNE) was amplified using primer HY17F ggtcgatgacagcctgcacgcgggcgac and HY17R aaagatcttacggcgcggtgatctcgttcg from the pv*RNE(ΔDBS)-YFP* GentR. The HY16F caccgcgccgtaagatcttttctacggggtctg and HY16R cgtgcaggctgtcatcgaccacgaccgac primers were used to amplify a pNTPs vector containing 1KB regions upstream and downstream of RNE’s NTD and a spectinomycin (Spec) cassette from the RNEΔCTD SpecR pNTPS138 plasmid [Ortiz-Rodriguez et al Nature Microbiology]. After running the DNA fragments on a 1% agarose gel, a GeneJet Gel Extraction kit was used to extract the PCR products. The GeneJET PCR Purification Kit was then used to purify the vector sample after it had been treated with Dpn1. The CTDΔDBS insert region was then assembled into the RNEΔCTD SpecR pNTPS138 vector via Gibson Assembly (NEB). The Gibson mix was then used to transform chemically competent *Escherichia coli* (E. coli) DH10B cells that were then selected on LB agar plates supplemented with Kanamycin (Kan) (30 ug/mL). Kan resistant colonies were screened for the presence of the CTDΔDBS insert region by PCR using the HY18F taagatcttttctacggggtctg and HY18R tcttcttcctcgtcgtcg primers and verified by Sanger sequencing (Genewiz). For the RNEΔDBS strain, the purified RNEΔDBS SpecR pNTPS138 (pHY001) plasmid was mated into *JS769* cells [Ortiz-Rodriguez et al bioRxiv] via tri-parental mating and plated on PYE agar plates supplemented with Nalidixic acid (Nal) + Spec + Kan (100 ug/mL Spec; 20 ug/mL Nal; 25 ug/mL Kan). The resulting kanR/specR resistant colonies were then inoculated into plain PYE and then counter-selected on PYE plates supplemented with Spec + 3% sucrose. Colonies that were resistant to Spec and sensitive to Kan were then identified and screened by PCR using primers Screen F tcggtcggcctctatatcc and Screen R gcgaaccctgaccaatctaa for the deletion of the DBS regions.

### JS495 VanA:: (double cys mutant(CTD)-YFPC-4) GentR

The RNE DNA fragment with the double cysteine-to-alanine mutations was synthesized (CATATGTCGAAGAAGATGCTGATCGACGCAGCACACGCCGAAGAGACGCGTGTGGTCGTCGTGGACGGTACCCGGGTTGAAGAGTTCGATTTCGAGAGCCAAACCCGCAAACAGCTTCGTGGAAACATCTATCTCGCCAAGGTGACGCGCGTTGAGCCCAGCCTCCAGGCTGCGTTCATCGAGTACGGCGGCAACCGTCACGGTTTCCTGGCGTTCAACGAGATCCACCCCGACTACTACCAGATCCCGGTCGCCGACCGCGAAGCGCTGATGCGCGACGACTCCGGTGACGACGAGGACGACACCCCGATCTCGCGTCGCGCCTCCGGCGGCGACGACGAAGACGACGTCAATGGCGGCGACCGCGCGGTCGACGATGATGACGATGACGTCGAAGAAGAACTGGCGCGCCGCAAGCGCCGCCTGATGCGCAAGTACAAGATCCAGGAAGTGATCCGCCGCCGGCAGATCATGCTGGTTCAGGTGGTCAAGGAAGAGCGTGGCAACAAGGGCGCGGCCCTGACCACCTATCTGTCGCTGGCCGGCCGCTACGGCGTCCTGATGCCCAACACCGCCCGTGGCGGCGGCATCAGCCGCAAGATCACGGCGGTGACTGACCGCAAGCGCCTGAAGAGCGTCGTCCAAAGCCTGGACGTGCCGCAAGGCATGGGTCTGATTGTCCGCACAGCCGGCGCCAAGCGCACCAAGGCCGAGATCAAGCGCGACTATGAGTACCTGCTGCGTCTGTGGGAGAACATCCGCGAGAACACGCTGCACTCGATCGCGCCGGCGCTGATCTACGAGGAAGAAGACCTCGTCAAACGCGCCATCCGCGACATGTACGACAAGGACCTGGACGGCATCTGGGTCGAGGGCGACGCCGGCTACAAGGAAGCGCGCGACTTCATGCGCATGCTAATGCCGAGCCAGGCCAAGAAGGTCTTCAACTACCGCGACCCGACCCCGCTGTTCGTGAAGAACAAGATCGAGGACCATCTGGCCCAGATCTATTCGCCGGTCGTTCCGCTGCGCTCGGGCGGCTATCTGGTGATCAACCAGACCGAGGCCCTGGTCGCCATCGACGTCAACTCGGGTAAGGCCACGCGCGAGCGCAACATCGAGGCCACCGCGCTGAAGACCAACTGCGAAGCGGCCGAGGAAGCCGCCCGTCAGCTGCGTCTGCGCGACCTGGCCGGCCTGATCGTCATCGACTTCATCGACATGGATGAAGCCAAGAACAACCGCACGGTCGAGAAGGTCCTGAAGGACGCGCTCAAGGACGACCGCGCGCGCATCCAGATGGGCAAGATCTCGGGCTTTGGCCTGATGGAGATCAGCCGTCAGCGTCGCCGCACCGGCGTGCTGGAAGGCACCACCCATGTCGCCGAACACGCCGAAGGCACCGGCCGTGTCCGTTCGGTGGAATCCAGCGCCCTGGCCGCCCTGCGCGCCGTCGAGGCCGAGGCCCTGAAGGGCTCGGGCAGCGTGATCCTGAAGGTCTCGCGCTCGGTCGGCCTCTATATCCTCAACGAAAAGCGCGATTATCTGCAGCGTCTGCTGACCACGCACGGCCTGTTCGTGTCGGTCGTGGTCGATGACAGCCTGCACGCGGGCGACCAGGAGATCGAGCGCACCGAGCTGGGCGAACGCATCGCCGTGGCCCCGCCGCCCTTCGTCGAGGAAGACGACGACTTCGATCCGAACGCCTACGACGACGAGGAAGAAGAAGACGACGTCATTCTCGATGACGAGGACGACACCGACCGCGAGGACACCGACGACGACGATGCGACGACGCGCAAGTCGGCGCGTGATGACGAGCGCGGCGACCGCAAGGGCCGTCGCGGGCGTCGCGACCGCAACCGCGGCCGCGGGCGTCGCGACGAGCGGGATGGCGAGACCGAGTCCGAGGACGAGGACGTCGTGGCCGAAGGCGCGGACGAGGATCGCGGCGAGTTTGGCGATGATGATGAAGGCGGTCGTCGCCGCCGTCGCCGGGGTCGTCGTGGCGGCCGTCGTGGCGGGCGCGAGGACGGCGATCGTCCGACCGACGCCTTCGTCTGGATCCGTCCGCGGGTGCCCTTCGGCGAGAACGTCTTCACCTGGCATGATCCGGCTGCGCTGGTCGGCGGTGGCGAGTCGCGTCGTCAGGCGCCCGAGCCGCGCGTCGATGCCGCTACCGAGGCCGCGCCGCGTCCCGAGCGGGCCGAGCGCGAAGAGCGCCCTGGCCGTGAACGTGGCCGTCGGGGTCGTGACCGGGGCCGTCGCCAGCGCGACGAGGCGCCGGTCGCCGAGATGACCTCGGTGGAAAGCGCGACTGTCGAGGCTGCGGAGCCGTTCGAGGCCCCCATCCTGGCGCCGCCGGTAATCGCCGG GCCGCCGGCCGACGTTTGGGTCGAACTGCCGGAAGTCGAGGAAGCGCCCAAGAAGCCCAAGCGCTCAAGGGCGCGCGGCAAGAAAGCGACTGAAACGTCCGTCGAAGCGATCGACACCGTCACCGAAGTCGCGGCGGAGGCTCCCGCCCCCGAGACCGCTGAACCCGAAGCCGTCGAGGTCGCTCCGCCGGCCCCCACGGTCGAGGCTGCGCCTGAGCCGGGACCGGTCGTCGAAGCCGTCGAGGAGGCCCAACCGGCCGAGCCGGATCCGAACGAGATCACCGCGCCGCCCGAAAAGCCCCGTCGGGGCTGGTGGCGCCGGGCGAATTC) and subcloned into pVYFPC-4 by GenScript. The resulting RNE(Cys-to-ala)-pVYFPC-4 plasmid was transformed into NA1000 cells and selected on PYE plates supplemented with Gentamycin (Gent) (5 ug/mL). The resulting GentR colonies were grown in liquid PYE + Gent (0.5 ug/mL).

### JS802 rne::pXRNEssrAC VanA:: (double cys mutant(CTD)-YFPC-4) KanR GentR

Phage lysate from strain *JS8* harboring RNE KanR under the xylose promoter was transduced into JS495 cells and selected on PYE + Gent + Kan + Xyl plates. The resulting GentR KanR colonies were inoculated into liquid PYE supplemented with Gent + Kan + Xyl.

### *JS376* rne::rne*ΔCTD*-eYFP gentR

525bp of the *Agrobacterium* RNase E gene up to the codon 692 was PCR amplified from the *A. tumefaciens* genome using primers agro_ntd-R gcgtaacgttcgaattcgcatcgtcttcttcttcaacgaagctcgg and agro_ntd-F cgtccaattgcatatgcttgctggtctcgttgtcatcg. The PCR product was digested with NdeI and EcoRI and ligated into pYFPC-4 [thanbichler ref] and verified by sangar sequencing. The resulting plasmid was then electroporated into *A. tumefaciens* as in [Al-husini mol cell 2018 ref].

## Acknowledgements

National Institute of Health. R01GM136863 to W.S.C., R35GM124733 to J.M.S. We’d also like to thank Huaiying Zhang for providing access for FRAP experiments. Wade Schnorr, Hannah Hunter and Kathryn Dzurik was supported by a US Department of Education GAANN grant, award number: P200A240158

