## Supplementary Materials for "BR-Bodies Facilitate Adaptive Responses and Survival During Copper Stress in *Caulobacter crescentus*"

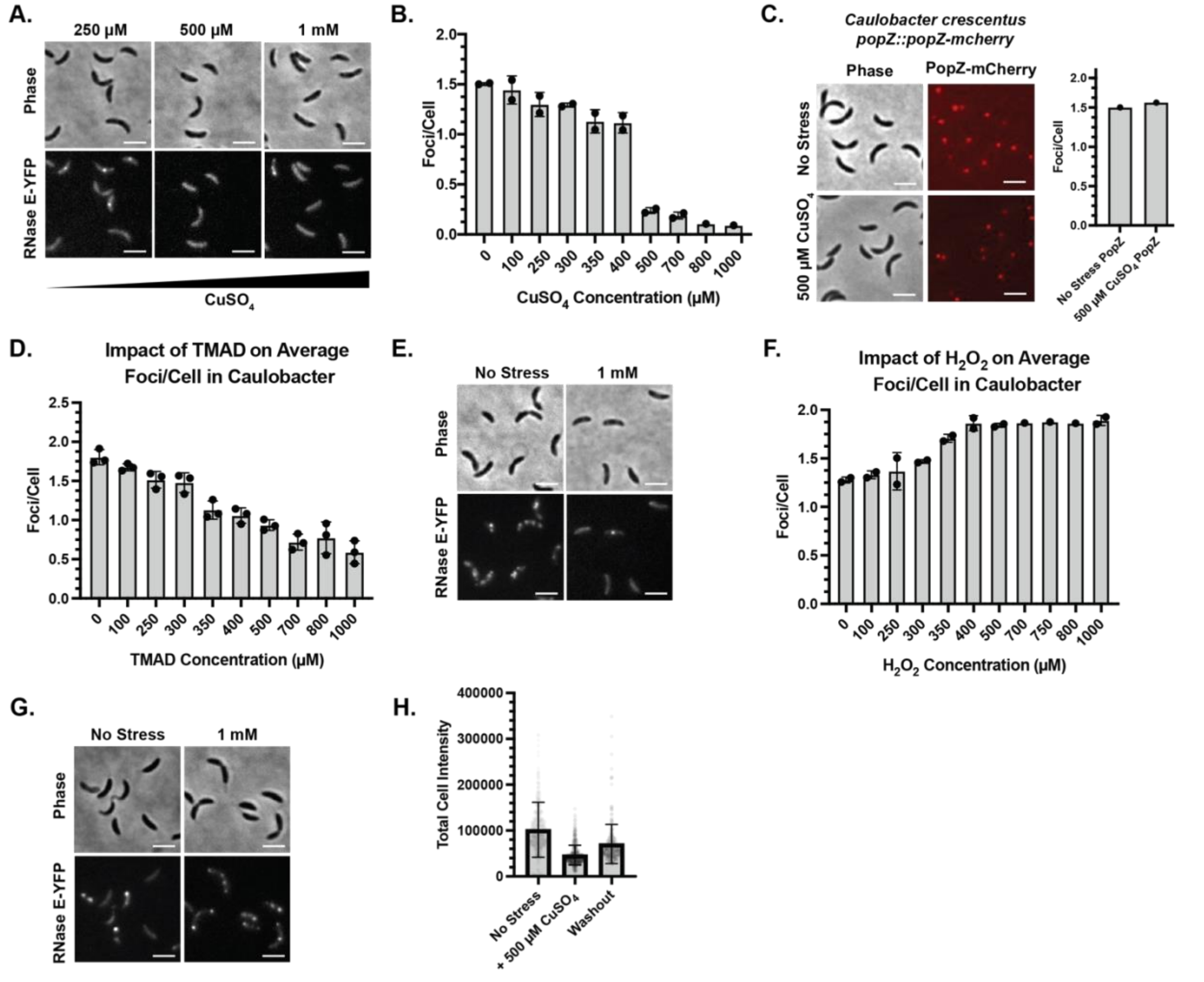


**SI Figure 1: Extensive *Caulobacter crescentus* BR-body dissolution is RNase E- and copper-specific with mild dissolution effects in the presence of TMAD and an increase in BR-bodies/cell in the presence of $H_2O_2$** (A) Representative phase contrast and fluorescence microscopy imaging of *C. crescentus* expressing RNase E-eYFP from its endogenous promoter in the presence of 250 µM $CuSO_4$, 500 µM $CuSO_4$, and 1 mM $CuSO_4$ for 8 min. BR-body dissolution is promoted with increasing concentrations of $CuSO_4$. The scale bar denotes 3 µm. (B) Quantification of the number of BR-bodies per cell in *C. crescentus* expressing RNase E-eYFP with varying concentrations of $CuSO_4$ from 0-1000 µM. (C) Phase contrast and fluorescence microscopy imaging of *C. crescentus* expressing PopZ-mCherry from its endogenous promoter in the absence and presence of 500 µM $CuSO_4$. Quantitative analysis of *C. crescentus* expressing PopZ-mCherry from its endogenous promoter in the absence and presence of 500 µM $CuSO_4$ (n=322,99 respectively). (D) Quantification of the number of BR-bodies per cell in *C. crescentus* expressing RNase E-eYFP with varying concentrations of TMAD from 0-1000 µM. (E) Representative phase contrast and fluorescence microscopy imaging of *C. crescentus* expressing RNase E-eYFP from its endogenous promoter in the absence and presence of 1 mM TMAD for 8 min. BR-body dissolution is promoted with increasing concentrations of TMAD. The scale bar denotes 3 µm. (F) Quantification of the number of BR-bodies per cell in *C. crescentus* expressing RNase E-eYFP with varying concentrations of $H_2O_2$ from 0-1000 µM. (G) Representative phase contrast and fluorescence microscopy imaging of *C. crescentus* expressing RNase E-eYFP from its endogenous promoter in the absence and presence of 1 mM $H_2O_2$ for 8 min. An increase in BR-bodies/cell is promoted with increasing concentrations of $H_2O_2$. The scale bar denotes 3 µm. (H) Quantification of total cell intensity in arbitrary units corresponding to $CuSO_4$ washout experiment with *C. crescentus* expressing RNase E-eYFP from its endogenous promoter (JS51).

**A.**

| Metals | PYE | EDTA | BCS | Nc | TTM |
|---|---|---|---|---|---|
| $Mg^{2+}$ | - | + | - | - | - |
| $Cu^{1+}$ | - | - | + | + | - |
| $Cu^{2+}$ | - | + | - | - | + |
| $Fe^{2+}$ | - | + | - | + | - |
| $Mn^{2+}$ | - | + | - | - | - |
| $Ni^{2+}$ | - | + | - | - | - |
| $Zn^{2+}$ | - | + | - | + | - |

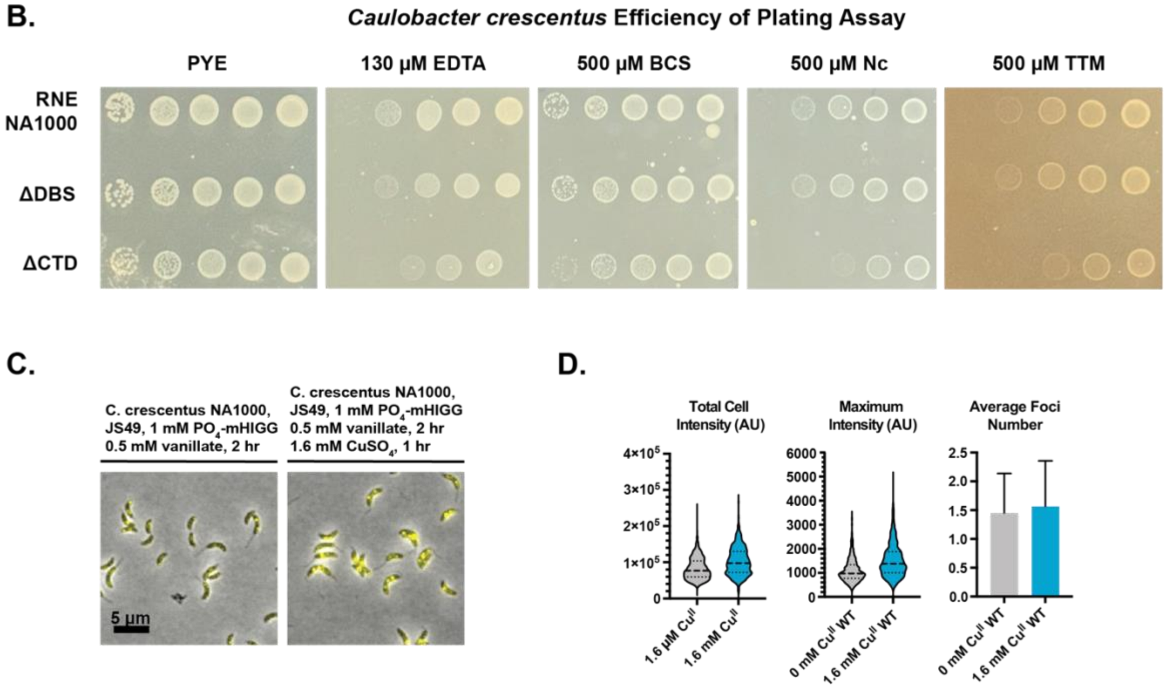


**SI Figure 2: Broad spectrum and copper-specific chelators promote an increase in RNase E foci intensity and confirm the importance of phase separation for high fitness under metal chelation** (A) Metal binding screen of broad spectrum EDTA metal chelator versus Bathocuproine sulfonate (BCS), Neocuproine (Nc) and Ammonium Tetrathiomolybdate (TTM). (B) Efficiency of Plating assay with wildtype *C. crescentus* RNase E, degradosome binding site mutant (ΔDBS) RNase E and NTD C-terminal deletion mutant (ΔCTD) RNase E in the absence and presence of EDTA and all copper chelators. BR-body phase separation provides enhanced fitness in the presence of broad-spectrum EDTA metal chelator and all copper-specific chelators. (C) Overlay of phase contrast and fluorescence microscopy imaging of a *C. crescentus* strain expressing the vanillate-inducible RNase E-eYFP (JS49, *vanA::rne-eYFP*) grown in modified HIGG media in the absence and presence of 1.6 mM $CuSO_4$ over 1 hour. (D) Quantification of total cell intensity in arbitrary units, maximum intensity in arbitrary units, and the average number of BR-bodies per cell in the absence (gray) and presence (blue) of 1.6 mM $CuSO_4$, respectively. BR-bodies were significantly more intense in cells grown under high Cu stress (high Cu: 1509±675 AU, n=1229; low Cu: 1097±472 AU, n=893, $p < 0.0001$). BR-body number was also significantly higher under high Cu stress (high Cu: 1.56±0.79, n=571; low Cu: 1.49±0.68, n=848, $p = 0.0038$). However, total cell intensity was also significantly different (high Cu 102350±38099 AU vs low Cu 82647±29702 AU, $p = 0.0027$).

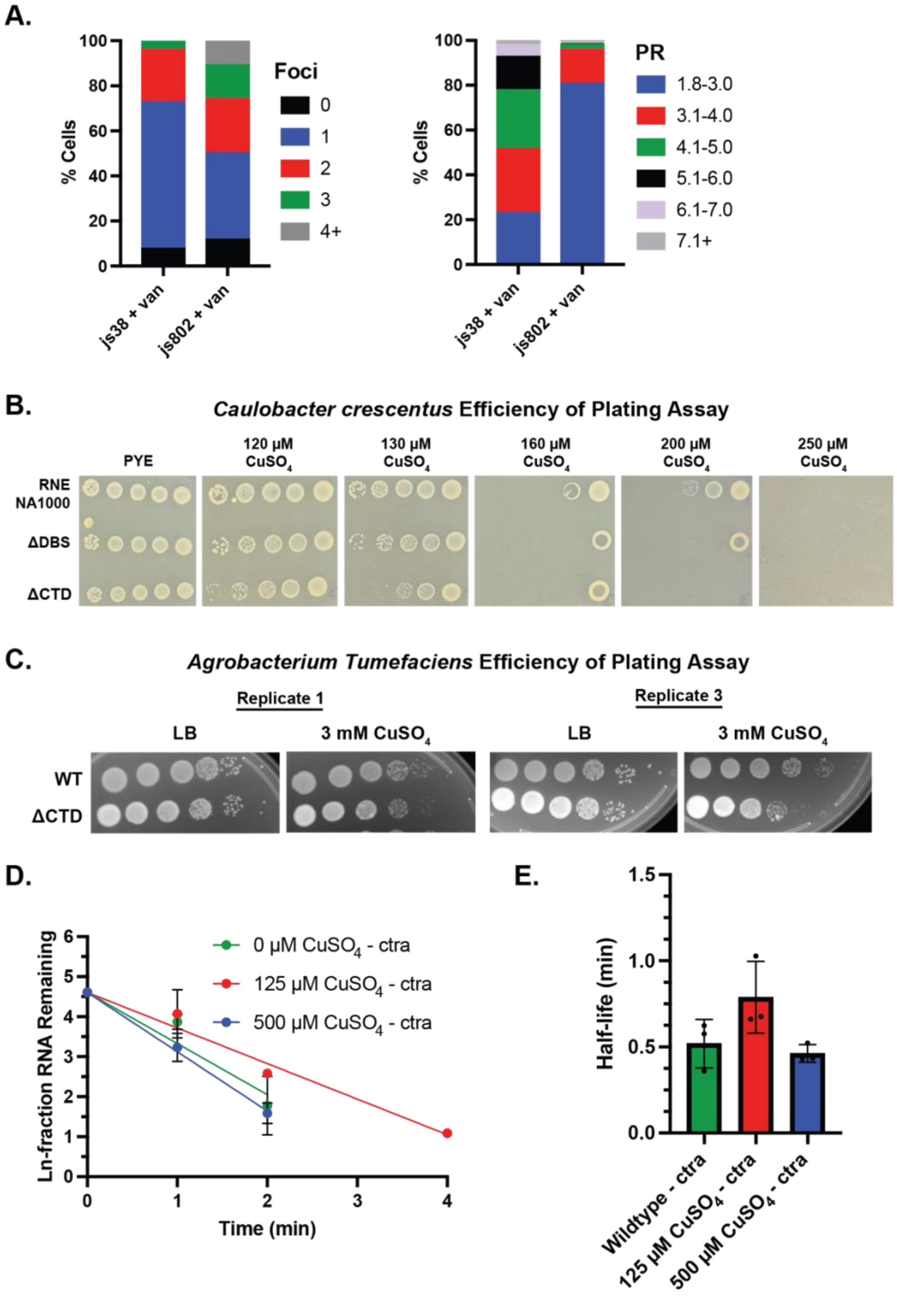


**SI Figure 3: Recruitment of RNase E clients into BR-bodies and BR-body phase separation provide enhanced fitness in the presence of increasing copper stress.** (A) Quantification of the number of BR-bodies per cell and partition ratio for wildtype (js38 + van) and mutant RNase E C461A, C464A (js802 + van). The RNase E mutant variant displays more heterogeneous foci/cell. (B) Efficiency of Plating assay with wildtype *C. crescentus* RNase E, degradosome binding site mutant (ΔDBS) RNase E and NTD C-terminal deletion mutant (ΔCTD) RNase E in the absence and presence of 120 μM $CuSO_4$, 130 μM $CuSO_4$, 160 μM $CuSO_4$, 200 μM $CuSO_4$ and 250 μM $CuSO_4$. (C) Efficiency of Plating assay with wildtype A*. tumefaciens C58* RNase E and the RNase e NTD C-terminal deletion mutant (ΔCTD) RNase E in the absence and presence of 3 mM $CuSO_4$. (D) Plot of Ln-fraction of RNA remaining for wild-type RNase E in the absence and presence of 125 μM $CuSO_4$ and 500 μM $CuSO_4$. (E) Quantification of mRNA half-life for wild-type RNase E in the absence and presence of 125 μM $CuSO_4$ and 500 μM $CuSO_4$ mRNA half-life decreases at elevated $CuSO_4$ concentrations.

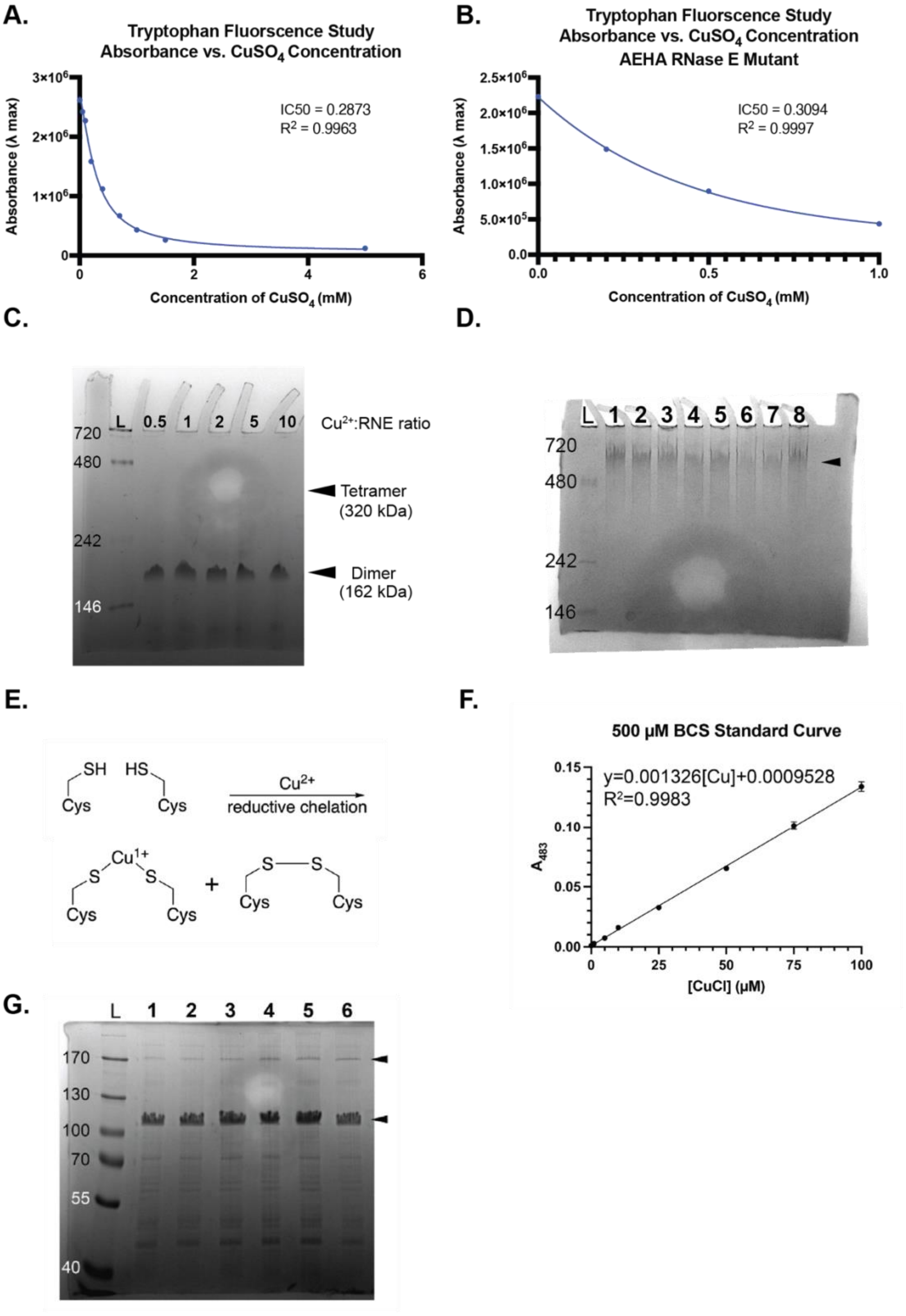

A.
Tryptophan Fluorscence Study
Absorbance vs. CuSO4 Concentration
IC50 = 0.2873
R2 = 0.9963
Absorbance (λ max)
Concentration of CuSO4 (mM)
B.
Tryptophan Fluorscence Study
Absorbance vs. CuSO4 Concentration
AEHA RNase E Mutant
IC50 = 0.3094
R2 = 0.9997
Absorbance (λ max)
Concentration of CuSO4 (mM)
C.
L 0.5 1 2 5 10
Cu2+:RNE ratio
720
480
242
146
Tetramer (320 kDa)
Dimer (162 kDa)
D.
L 1 2 3 4 5 6 7 8
720
480
242
146
E.
SH HS
Cys Cys
Cu2+
reductive chelation
Cu1+
S S
Cys Cys
+
S S
Cys Cys
F.
500 μM BCS Standard Curve
y=0.001326[Cu]+0.0009528
R2=0.9983
A483
[CuCl] (μM)
G.
L 1 2 3 4 5 6
170
130
100
70
55
40

**Figure S4:** (A) Absorbance vs. $CuSO_4$ concentration for unlabeled RNase E (451-898) (B) Absorbance vs. $CuSO_4$ concentration for C461A, C464A unlabeled RNase E (451-898) variant. Titration by Cu(II) monitored by protein intrinsic fluorescence intensity quenching. Increasing concentrations of $CuSO_4$ stress result in a decrease in tryptophan fluorescence intensity. (C) Native PAGE analysis of RNase E (451-898) upon $Cu^{2+}$ incubation. Oligomerization analysis of RNase E (451-898) resolved on an 8% native polyacrylamide gel. L: resolved NativeMark (Thermofisher) in kDa. Lanes 1-5: Analysis of 6 μM RNase E (451-898) incubated with (1) no metals, (2) 3 μM $CuSO_{44}$, (3) 6 μM $CuSO_4$, (4) 30 μM $CuSO_4$, (5) 60 μM $CuSO_4$. Triangles represent apparent molecular weights of an RNase E (451-898) dimer (*bottom*, 200 kDa) and predicted RNase E tetramer based on the dimer mass (*top*, 400 kDa). (D) Native PAGE analysis of RNase E (1-898, D403C) after $Cu^{2+}$ incubation. Oligomerization analysis of RNase E (1-898, D403C) resolved on an 8% native polyacrylamide gel. L: resolved NativeMark (Thermofisher) in kDa. Lanes 1-8: Analysis of 6 μM RNase E (1-898, D403C) incubated with (1) no metals, (2) 0.5X $CuSO_4$, (3) 1X $CuSO_4$, (4) 2.5X $CuSO_4$, (5) 5X $CuSO_4$, (6) 10X $CuSO_4$, (7) 10X $ZnSO_4$, (8) 100X $ZnSO_4$. Triangle represents the apparent mass of the protein dimer (625 kDa). (E) Reductive chelation of $Cu^{2+}$ by proximal cysteines. The proximal cysteines in RNase E could be oxidized by $Cu^{2+}$ by reductive chelation. $Cu^{2+}$ can interact with proximal RNase E cysteines, oxidizing one set of cysteines to cysteine and chelating the other as $(Cys)_2Cu^{1+}$. (F) Bathocuproine sulfonate standard curve with CuCl. Various concentrations of CuCl dissolved in 100 mM Tris-Cl pH 8.0 with 1 mM DTT was allowed to react with 500 μM BCS for 2 min prior to $A_{483}$ measurement. The extinction coefficient was determined to be 13260 $M^{-1}cm^{-1}$ based on the slope of a fit linear regression. Error was based on three replicates. (G) Non-reducing SDS-PAGE analysis of RNase E (451-898) upon $Cu^{2+}$ incubation. Oligomerization analysis of RNase E (451-898) resolved on a 10% non-reducing SDS polyacrylamide gel. L: resolved PageRuler (Thermofisher) in kDa. Lanes 1-5: Analysis of 6 μM RNase E (451-898) incubated with (1) no metals, (2) 0.25X $CuSO_4$, (3) 0.5X $CuSO_4$, (4) 1X $CuSO_4$, (5) 5X $CuSO_4$, (6) 10X $CuSO_4$. Triangles represent apparent molecular weights of an RNase E (451-898) dimer (bottom, 110 kDa) and predicted RNase E tetramer based on the dimer mass (top, 220 kDa).

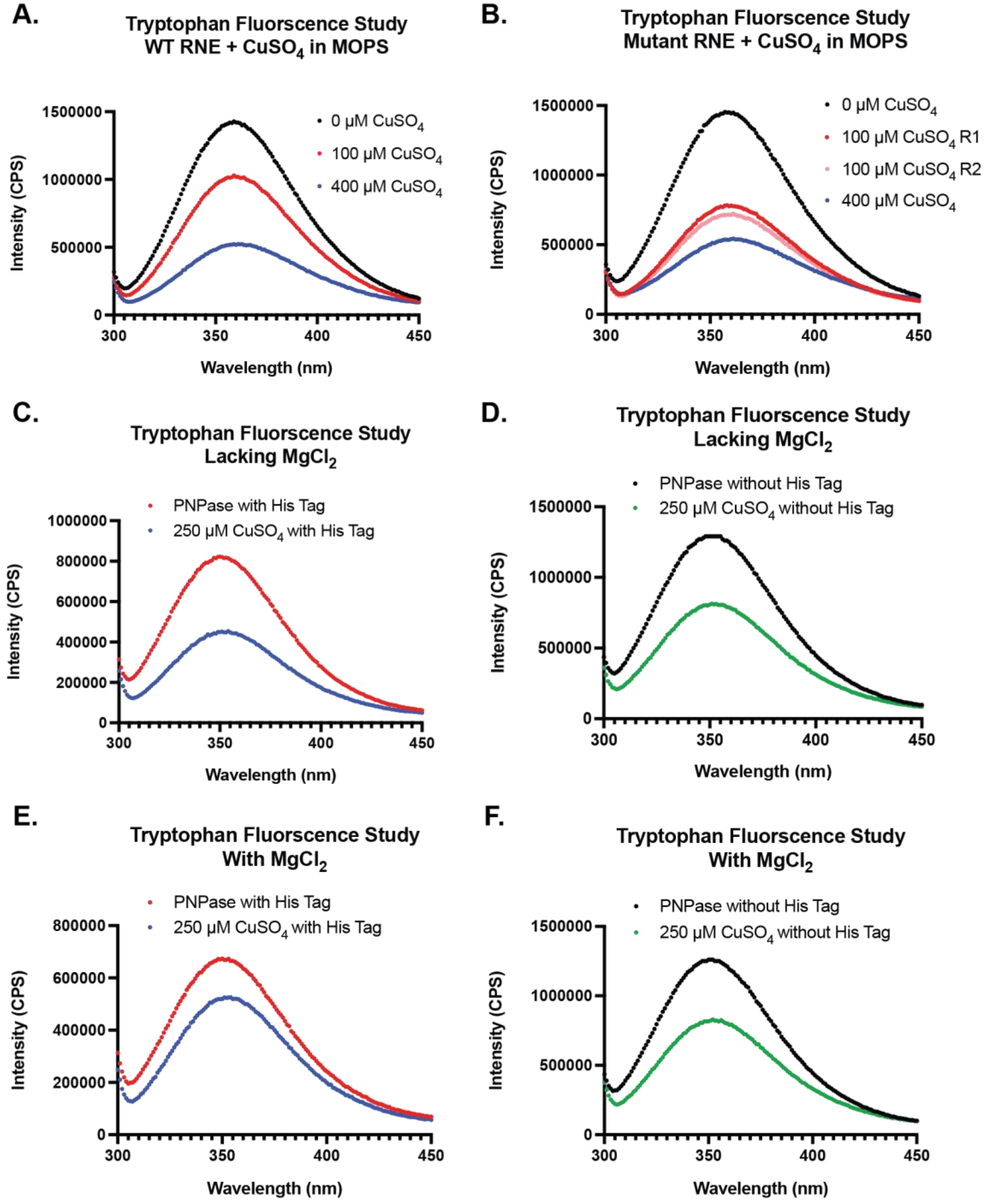
A.
Tryptophan Fluorscence Study
WT RNE + CuSO4 in MOPS
0 μM CuSO4
100 μM CuSO4
400 μM CuSO4
Intensity (CPS)
Wavelength (nm)
B.
Tryptophan Fluorscence Study
Mutant RNE + CuSO4 in MOPS
0 μM CuSO4
100 μM CuSO4 R1
100 μM CuSO4 R2
400 μM CuSO4
Intensity (CPS)
Wavelength (nm)
C.
Tryptophan Fluorscence Study
Lacking MgCl2
PNPase with His Tag
250 μM CuSO4 with His Tag
Intensity (CPS)
Wavelength (nm)
D.
Tryptophan Fluorscence Study
Lacking MgCl2
PNPase without His Tag
250 μM CuSO4 without His Tag
Intensity (CPS)
Wavelength (nm)
E.
Tryptophan Fluorscence Study
With MgCl2
PNPase with His Tag
250 μM CuSO4 with His Tag
Intensity (CPS)
Wavelength (nm)
F.
Tryptophan Fluorscence Study
With MgCl2
PNPase without His Tag
250 μM CuSO4 without His Tag
Intensity (CPS)
Wavelength (nm)

**Figure S5:** (A) Unlabeled wildtype RNase E (451-898) titration by Cu(II) in MOPS buffer monitored by protein intrinsic fluorescence intensity quenching. Increasing concentrations of $CuSO_4$ stress result in a decrease in tryptophan fluorescence intensity. (B) Unlabeled mutant RNase E C461A/C464A titration by Cu(II) in MOPS buffer monitored by protein intrinsic fluorescence intensity quenching. Increasing concentrations of $CuSO_4$ stress result in a decrease in tryptophan fluorescence intensity. (C) Unlabeled PNPase with a histidine tag titration by Cu(II) in the absence of $MgCl_2$ monitored by protein intrinsic fluorescence intensity quenching. Increasing concentrations of $CuSO_4$ stress result in a decrease in tryptophan fluorescence intensity. (D) Unlabeled PNPase without a histidine tag titration by Cu(II) in the absence of $MgCl_2$ monitored by protein intrinsic fluorescence intensity quenching. Increasing concentrations of $CuSO_4$ stress result in a decrease in tryptophan fluorescence intensity. (E) Unlabeled PNPase with a histidine tag titration by Cu(II) in the presence of $MgCl_2$ monitored by protein intrinsic fluorescence intensity quenching. Increasing concentrations of $CuSO_4$ stress result in a decrease in tryptophan fluorescence intensity. (F) Unlabeled PNPase without a histidine tag titration by Cu(II) in the presence of $MgCl_2$ monitored by protein intrinsic fluorescence intensity quenching. Increasing concentrations of $CuSO_4$ stress result in a decrease in tryptophan fluorescence intensity.

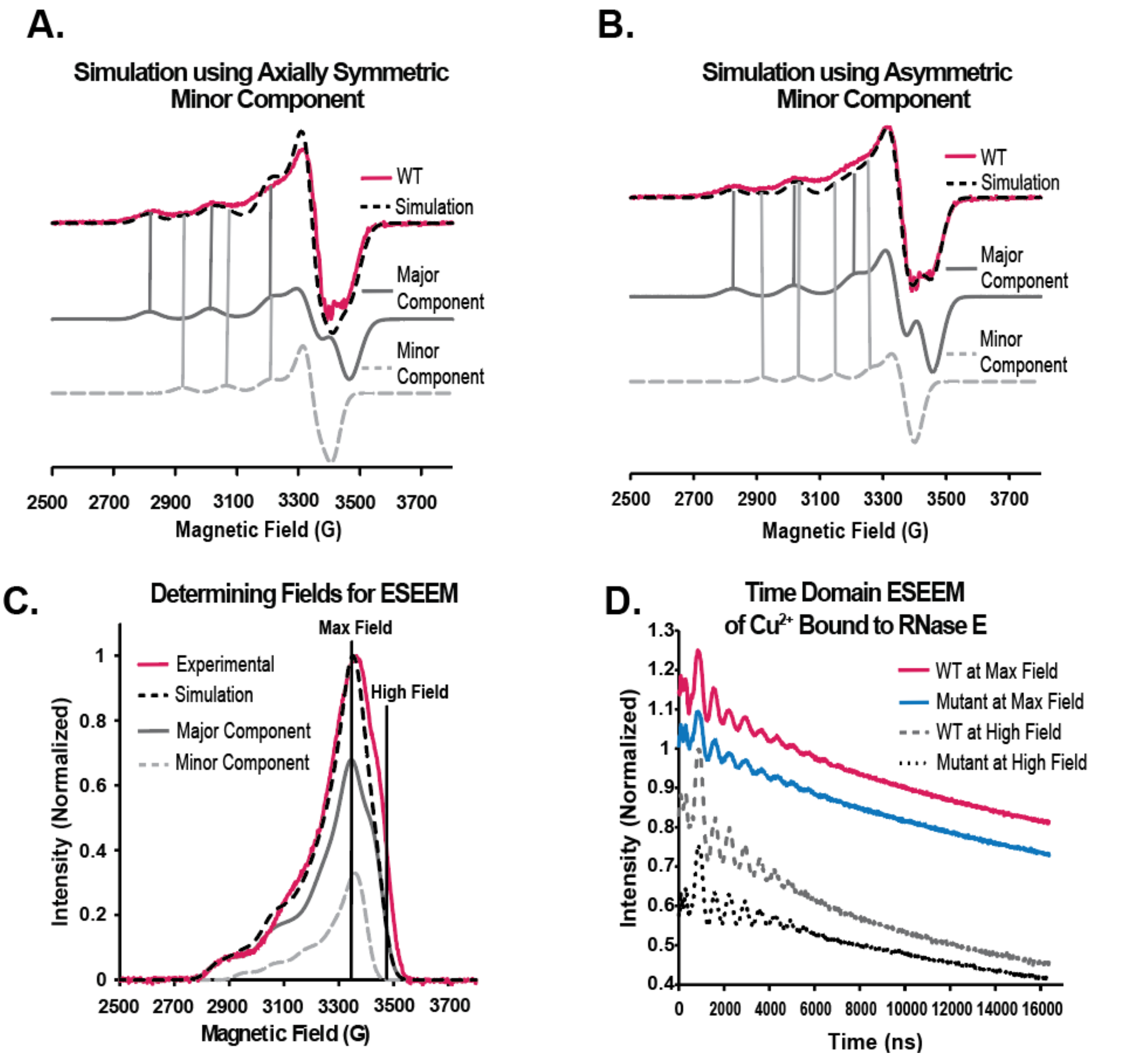


**Figure S6.** Simulation of RNase E wild-type (pink) is overlaid in dashed black. The RNase E spectrum shows two components. The major component (grey) accounts for 75% of the simulation (dashed black), and the minor component (dashed light grey) accounts for the remaining 25% of the simulation (dashed black). (A) Simulation of RNase E wild-type fit with an axially symmetric minor component (light grey). (B) Simulation of RNase E wild-type fit with an asymmetric minor component (light grey). (C) Field swept spectrum of the RNase E wild-type (pink) overlaid with the integrated CW-EPR simulation (dashed black) using an asymmetric coordination geometry for the minor component (dashed light grey). ESEEM experiments were carried out at the field with the maximum intensity (3340 G) and a higher field (3480 G), highlighted with black lines. At the high field, only the major component (grey) contributes to the spectrum. (D) Raw time domain ESEEM signal collected at the maximum intensity and higher field. Data has been normalized and offset for clarity.

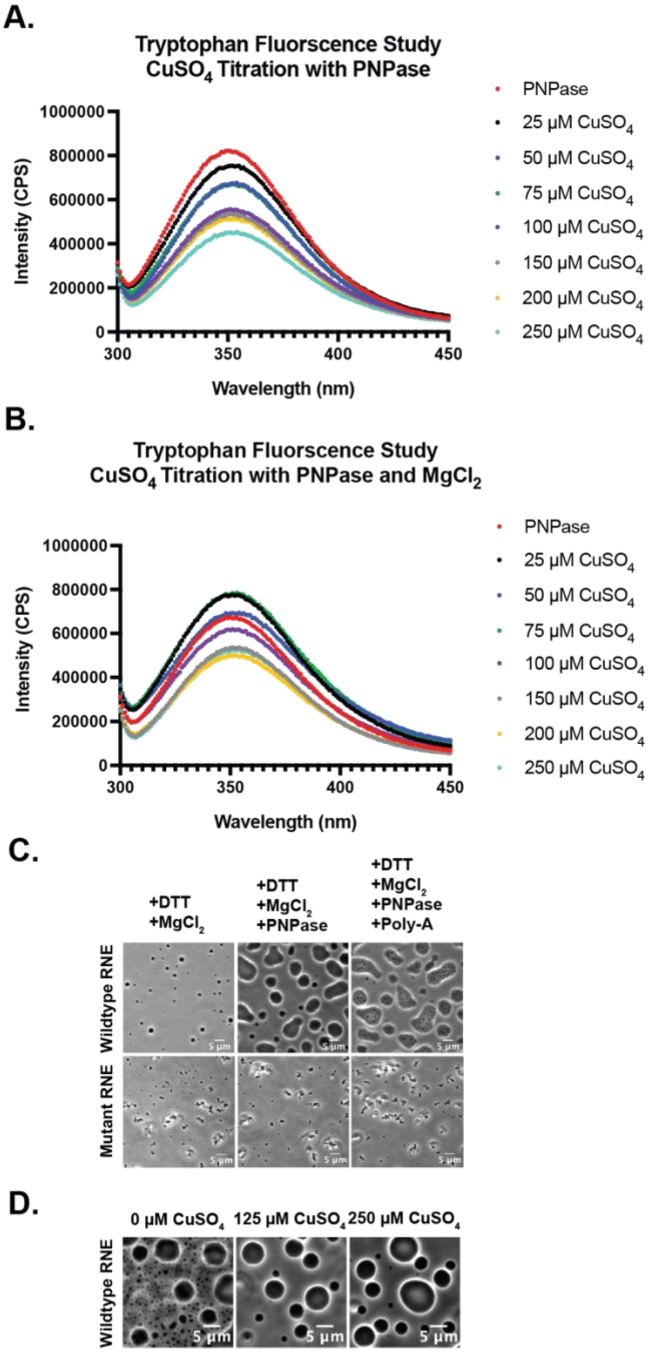


**Figure S7:** (A) Unlabeled PNPase titration by Cu(II) monitored by protein intrinsic fluorescence intensity quenching. Increasing concentrations of $CuSO_4$ stress result in a decrease in tryptophan fluorescence intensity. (B) Unlabeled PNPase titration by Cu(II) in the presence of $MgCl_2$ monitored by protein intrinsic fluorescence intensity quenching. Increasing concentrations of $CuSO_4$ stress result in a decrease in tryptophan fluorescence intensity. (C) The addition of PNPase increases RNase E droplet size, enhancing RNase E's ability to sequester Poly-A into RNase E condensates, as can be visualized by the presence of dark speckles within RNase E-PNPase droplets. In comparison, in the presence of PNPase and poly(A), the RNase E (451-898) C461A/C464A variant forms aggregate-like assemblies. Samples were incubated for 1 hour prior to imaging. (D) RNase E droplets maintain spherical morphology in the presence of PNPase with increasing $CuSO_4$ concentration.

**Table 1.** A summary of g and hyperfine tensors for $Cu^{2+}$ in RNase E samples was obtained via simulations using EasySpin.

| Sample | Major Component | | | | Axially Symmetric Minor Component | | | |
|---|---|---|---|---|---|---|---|---|
| | $g_\parallel$ | $A_\parallel$ (G) | $g_\perp$ | $A_\perp$ (G) | $g_\parallel$ | $A_\parallel$ (G) | $g_\perp$ | $A_\perp$ (G) |
| WT | 2.218 | 187 | 2.058 | 3 | 2.203 | 136 | 2.062 | 3 |
| Mutant | 2.218 | 187 | 2.058 | 3 | 2.203 | 136 | 2.062 | 3 |
| $CuSO_4$ in HEPES | 2.270 | 161 | 2.053 | 5 | N/A | | | |

| | Major Component | | | | Asymmetric Minor Component | | | | | |
|---|---|---|---|---|---|---|---|---|---|---|
| | $g_\parallel$ | $A_\parallel$ (G) | $g_\perp$ | $A_\perp$ (G) | $g_{zz}$ | $A_{zz}$ (G) | $g_{yy}$ | $A_{yy}$ (G) | $g_{xx}$ | $A_{xx}$ (G) |
| WT | 2.218 | 187 | 2.058 | 3 | 2.233 | 112 | 2.058 | 7 | 2.040 | 11 |
| Mutant | 2.218 | 187 | 2.058 | 3 | 2.233 | 112 | 2.058 | 7 | 2.040 | 11 |

**Table 2 Plasmid Construction Table**

| Plasmid Name | Description | Vector | Digest Site | Template | Sense Primer | Antisense Primer |
|---|---|---|---|---|---|---|
| pDT058 | *pET28::rne*(451-898)-YFP | pET28 | -- | -- | -- | -- |
| pDT060 | pET28-RNase E(1-898) | pET28 | -- | -- | -- | -- |
| pDT074 | RNase E (451-898) | pTEV5 | NheI | pDT058 | DT061 | DT085 |
| pDT124 | RNase E (451-898, C461A, C464A)-eYFP | pTEV5 | NheI | pDT058 | DT061 | DT168 |
| | | | | pDT058 | DT167 | DT123 |
| pDT156 | RNase E (451-898, C464A)-eYFP | pTEV5 | NheI | pDT058 | DT061 | DT203 |
| | | | | pDT058 | DT202 | DT123 |
| pDT157 | RNase E (451-898, C461A)-eYFP | pTEV5 | NheI | pDT058 | DT061 | DT204 |
| | | | | pDT058 | DT205 | DT123 |
| pDT280 | MBP-RNase E (1-898, D403C) | pTEV5 | NheI | pTEV6 | DT471 | DT472 |
| | | | | pDT060 | DT423 | DT275 |
| | | | | pDT060 | DT274 | DT085 |
| pMJC0094 | CcPNPase | pTEV5 | NheI | -- | -- | -- |
| pHY001 | *pNTPS-rne::rneΔDBS specR* | pNTPS-138 | -- | pv*RNE(ΔDBS)-YFP* GentR, and pNTPS-*rne::rneΔCTD specR* | HY16F, HY17F | HY16R, HY17R |
| pVYFPRNE-doublecysala-YFP | PvanA::rne(1-898, C461A, C464A)-eYFP gentR | pvYFP-C4 | EcoR1, Nde1 | synthesized | -- | -- |
| patNTD-YFP | prne*ΔCTD*-eYFP gentR | pYFPC-4 | NdeI/EcoR1 | *A. tumefaciens* genome | agro_ntd-F | agro_ntd-R |

* For plasmid construction of strains by the Schrader Lab (JS51, JS38), please see Al-Husini *et al*. *Mol. Cell.* **2018**, *71*, 1027-1039.

**Table 3 Strain Collection**

| Strain Name | Description | Organism | Source |
|---|---|---|---|
| DH5α | Bacterial cloning strain | *E. coli* | Invitrogen |
| Top10 | Bacterial cloning strain | *E. coli* | Invitrogen |
| BL21 (DE3) | Bacterial expression strain | *E. coli* | Novagen |
| Rosetta™ (DE3) | Bacterial expression strain | *E. coli* | Novagen |
| NA1000 | Synchronizable version of CB15 | *C. crescentus* | Shapiro Lab |
| DTT008 | pDT058 | BL21 (DE3) | Dylan's Thesis/Childers Lab |
| DTT028 | pDT074 | Rosetta™ (DE3) | Dylan's Thesis/Childers Lab |
| JS38 | *vanA::rne(1-577)-YFP* | NA1000 | Al-Husini et al Mol Cell 2018 |
| JS51 | *rne::rne(1-898)-YFP* | NA1000 | Al-Husini et al Mol Cell 2018 |
| WSC1748 | ParB::CFP-ParB; PopZ::mcherry-PopZ | Caulobacter with plasmid | This work/Childers Lab |
| | | NA1000 | |
| DTT248 | PvanA::RNase E(1-898, C461A, C464A)-eYFP, Prne::pXrnessraC | *C. crescentus* | This work/Schrader Lab |
| JS802 | | NA1000 | |
| DTT249 | PvanA::RNase E(1-898)-eYFP, Prne::pXrnessraC | *C. crescentus* | Al-Husini et al Mol Cell 2018 |
| JS38 | | NA1000 | |
| JS495 | PvanA::RNase E(1-898, C461A, C464A)-eYFP | *C. crescentus* | This Work/Schrader Lab |
| MJC192 | PNPase | *C. crescentus* | This work/Childers Lab |
| JS5 | rne::rne-eYFP | *Agrobacterium tumefaciens* | Al-husini et al Mol cell 2018 |
| JS376 | rne::rneΔCTD-eYFP | *Agrobacterium tumefaciens* | This work/Schrader Lab |
| JS769 | rne::rneΔCTD | *C. crescentus* | Ortiz Rodriguez et al. |
| JS801 | rne::rneΔDBS | *C. crescentus* | This work/Schrader Lab |

### Table 4 Reagents

Reagents

| REAGENT or RESOURCE | SOURCE | IDENTIFIER |
|---|---|---|
| Chemicals, peptides, and recombinant proteins | | |
| Gibson Master Mix | New England BioLabs Inc. | E2611S |
| Copper (II) sulfate pentahydrate | Sigmaaldrich | 7758-99-8 |
| TRizol Reagent | 15596018 | Ambion |
| Spectinomycin | Sigmaaldrich | S6501-25G |
| Rifampicin | Sigmaaldrich | R7382-1G |
| Kanamycin | Sigmaaldrich | K1377-5G |
| Nalidixic Acid | Sigmaaldrich | N8878-5G |
| Sucrose | sigmaaldrich | S5016-25G |
| Phusion DNA polymerase | Thermo Scientific | F-530L |
| RNAprotect Bacterial Reagent | QIAGEN | 76506 |
| Luna® Universal One-Step RT-qPCR Kit | NEB | E3005L |
| Qubit RNA HS Assay Kit | Thermo Fischer Scientific | Q32851 |
| Chloroform | Thermofisher scientific | AC423550010 |
| Glycogen, RNA grade | Thermofisher scientific | RO551 |
| 2-PROPANOL, ANHYDROUS<br>ETHANOL-D6 (D, 99%), ANHYD.<br>EDTA (0.5 M), pH 8.0, RNase-free | Sigmaaldrich<br>Sigmaaldrich<br>Sigmaaldrich | 67-63-0<br>1516-08-1 |

| REAGENT or RESOURCE | SOURCE | IDENTIFIER |
|---|---|---|
| Tris (1 M), pH 7.0, RNase-free | Thermofisher scientific | AM9261 |
| Agar | Thermofisher scientific | AM9851 |
| Bactopeptone | Fisherchemicals | DF0001-17-0 |
| Yeast extract | Thermofisher scientific | 211677 |
| UltraPure™ Ethidium Bromide, 10 mg/mL | sigmaaldrich | 92144-500G-F<br>15585011 |
| Thermo Scientific™ TriTrack DNA Loading Dye (6X) | Thermofisher scientific | FERR1161 |
| Magnesium sulfate | Thermofisher scientific | M7506-1KG |
| Calcium chloride (97%) | sigmaaldrich | 746495-500G |
| Luria Broth Base | sigmaaldrich | |
| FD Dpn1 | sigmaaldrich | 12795084 |
| T4 DNA Ligase | Thermofisher scientific | ER1701 |
| Agarose | Thermofisher scientific | EL0011 |
| RNAprotect Bacteria Reagent | Qiagen | A7705 |
| Gentamycin Sulfate | Sigmaaldrich | 76506 |
| Vanillic acid | Sigmaaldrich | 345814-1GM |
| D(+)-Xylose, 99+% | Thermofisher scientific | H36001-25G<br>141005000 |

**Author contributions:**

H.Y. has generated *JS801* and *JS802* and conducted all mRNA half-life measurements.

J.G. has helped with the screening and generation of *JS801*. AG has generated strain JS495.
